# The 4.4 Å structure of the giant Melbournevirus virion belonging to the *Marseilleviridae* family

**DOI:** 10.1101/2021.07.14.452405

**Authors:** Raymond N. Burton-Smith, Hemanth Kumar Narayana Reddy, Martin Svenda, Chantal Abergel, Kenta Okamoto, Kazuyoshi Murata

## Abstract

Members of *Marseilleviridae*, one family of icosahedral giant viruses classified in 2012 have been identified worldwide in all types of environments. The virion shows a characteristic internal membrane extrusion at the five-fold vertices of the capsid, but its structural details need to be elucidated. We now report the 4.4 Å cryo-electron microscopy structure of the Melbournevirus capsid. An atomic model of the major capsid protein (MCP) shows a unique cup structure on the trimer that accommodates additional proteins. A polyalanine model of the penton base protein shows internally extended N- and C-terminals, which indirectly connect to the internal membrane extrusion. The Marseilleviruses share the same orientational organisation of the MCPs as PBCV-1 and CroV, but do not appear to possess a protein akin to the “tape measure” of these viruses. Minor capsid proteins named PC-β, zipper, and scaffold are proposed to control the dimensions of the capsid during assembly.

## Introduction

Members of the *Marseilleviridae* family belonging to the nucleo-cytoplasmic large DNA viruses (NCLDVs) (Brandes and Linial, 2019) infect Acanthamoeba (Colson et al., 2013), and present a highly complex ∼360 kbp genome and icosahedral capsid with a diameter of ∼250 nm. The superfamily of NCLDVs was initially proposed by Iyer et al. (Iyer et al., 2001) and currently consist of nine families and several subfamilies of viruses (Subramaniam et al., 2020). According to ICTY classification, the members of the *Nucleocytoviricota* realm (PMID: 31187277) are restricted to seven families of giant viruses possessing double-stranded DNA and targeting a wide range of eukaryotic hosts. Since the first discovery of Marseillevirus in 2009 (Boyer et al., 2009), the *Marseilleviridae* has expanded to more than 10 members and classified into five lineages, A to E (Aoki et al., 2019; Fabre et al., 2017). Members of *Marseilleviridae* have been found in some human patients, but their link with a particular disease is yet to be identified (Aherfi et al., 2016; Kutikhin et al., 2014).

Melbournevirus is a member of the *Marseilleviridae* isolated from a freshwater pond in Melbourne, Australia in 2014 (Doutre et al., 2014). It shares common genomic and structural features with the other members of the family and classifies into lineage A. The first cryo-EM structure was reported at 26 Å resolution by using a 200 kV microscope (Okamoto et al., 2018), revealing the major capsid protein (MCP) array and a triangulation number of the icosahedral capsid as T=309. In addition, characteristic morphological features of Melbournevirus capsids were elucidated such as an extrusion of the internal membrane at the five-fold vertices of the capsid and a nucleoid containing a large density body, the nature of which has not yet been elucidated.

The sub-nanometer resolution structure of tokyovirus, one member of the *Marseilleviridae*, has been reported using high voltage cryo-electron microscopy (cryo-HVEM) (Chihara et al., 2021). The map did not reach a high enough resolution to be able to build *de novo* the structures of the various minor capsid proteins (mCPs) and only allowed a semi-fitted model for the MCP and a hypothetical fit of the PBCV-1 penton protein (Chihara et al., 2021). However, manual data collection with the HVEM microscope restricted the number of images collected, and the current resolution is limited to ∼8 Å after using a capsid-focussed mask. The tokyovirus cryo-EM map elucidated the novel capsid protein network of the giant virus including the scaffold proteins which could assist in the formation of the characteristic internal membrane extrusion at the five-fold vertices of the capsid and function in a similar manner to a tape measure protein (Xian et al., 2020). Furthermore, it suggested a glycosylated cap protein was sitting on the MCP trimer (Chihara et al., 2021).

High resolution cryo-EM maps of the giant viruses have been reported in three structures exceeding 5 Å resolution: 3.5 Å for PBCV-1 (Fang et al., 2019), 4.6 Å and 4.8 Å for African swine fever virus (ASFV) (Liu et al., 2019; Wang et al., 2019). These structures utilised 300 kV microscopes and were reconstructed with a technique termed “block based” reconstruction (Zhu et al., 2018) to improve obtained resolutions. Leveraging the high symmetry of icosahedral viruses, this technique can decrease the size of boxes necessary for 3D reconstruction and allow localized defocus refinement across the particle. For all objects >150 nm, this defocus gradient is considerable and has a negative impact on the attainable resolution. Consequently, they successfully constructed the atomic structures of the MCP and several mCPs in PBCV-1 and ASFV. In PBCV-1, the structures of 13 different types of mCP were identified in addition to the MCP (Fang et al. 2019), while in ASFV, the structures of four mCPs were determined in addition to the MCP (Liu et al., 2019; Wang et al., 2019).

Here, we report the structure of Melbournevirus to 4.9 Å for the whole particle, with only 3,124 particle images, and a maximum resolution of 4.42 Å obtained on the acquired dataset for the five-fold, three-fold and two-fold “blocks”, using symmetry expansion, recentring and defocus refinement. This is functionally the same as block-based reconstruction (Zhu et al., 2018) which has been applied to the study of other giant viruses but operates entirely within the RELION (Fernandez-Leiro and Scheres, 2017; Scheres, 2012; Zivanov et al., 2018; Zivanov et al., 2020) processing suite. The capsid, particularly in the block-based reconstructions, permits clearer views of the complex lattice of capsid proteins. While the hard limit of resolution in this data makes residue identification difficult, poly-alanine chain models could be created in some cases. An atomic model of the MCP, and a polyalanine model of the penton base protein which has no identified polypeptide sequence were built and compared with other giant viruses reported previously. The Marseilleviruses share the same orientational organisation of the major capsid proteins as PBCV-1 and CroV, but do not appear to possess a protein akin to the “tape measure” of these viruses, and the internal scaffold array does not extend to the pentasymmetron. This means some other mechanism to control the dimensions of the capsid during assembly, possibly utilising the PC-β, zippers, and scaffold proteins in concert, is required. These differences demonstrate the variety and biological flexibility of the giant viruses, while simultaneously utilising similar strategies for some critical features. The cryo-EM map of Melbournevirion provides clearer views of the complex lattice of the capsid, and the molecular interactions between capsid proteins, revealing the functional viral capsid network.

## Results

### The structure of Melbournevirus

The image processing flowchart and the dataset collection details of the whole-virion 3D reconstruction is shown in Fig. S1 and Table 1. Briefly, 3,367 micrograph movies of ice embedded Melbournevirions, in two independently collected datasets, consisting of 20 and 40 frames per micrograph respectively, were collected with a 300 kV Titan Krios (Thermo Fisher Scientific) microscope using automated image acquisition software. The image data were processed in the RELION 3.1 suite (Zivanov et al., 2018; Zivanov et al., 2020) throughout, as described in Methods. Using Ewald sphere correction, a 4.9Å global resolution map of Melbournevirus was reconstructed by assuming icosahedral symmetry (Fig. 1, Fig. S1F). Externally, Melbournevirus shares a high structural similarity to that of tokyovirus (Chihara et al., 2021), another member of the *Marseilleviridae*, sharing the same triangulation number (T=309), and the same MCP pattern on the capsid surface. Like tokyovirus, it shows a uniform MCP array each with a lower resolution “cap” density (Chihara et al., 2021). Internally, the inner membrane shows the characteristic extrusion at the five-fold axis, and the scaffold network between the capsid layer and inner membrane is comparable to that of tokyovirus (red circles in Fig. 1B). A low resolution “plug” of density fills a space between the pentasymmetron and the internal membrane extrusion (arrow in Fig. 1B), although it does not directly contact the internal membrane. These components may be flexible in the structure because they are not well resolved even in the non-symmetrized map of the block-based reconstruction (Fig. 2).

**Table 1.**
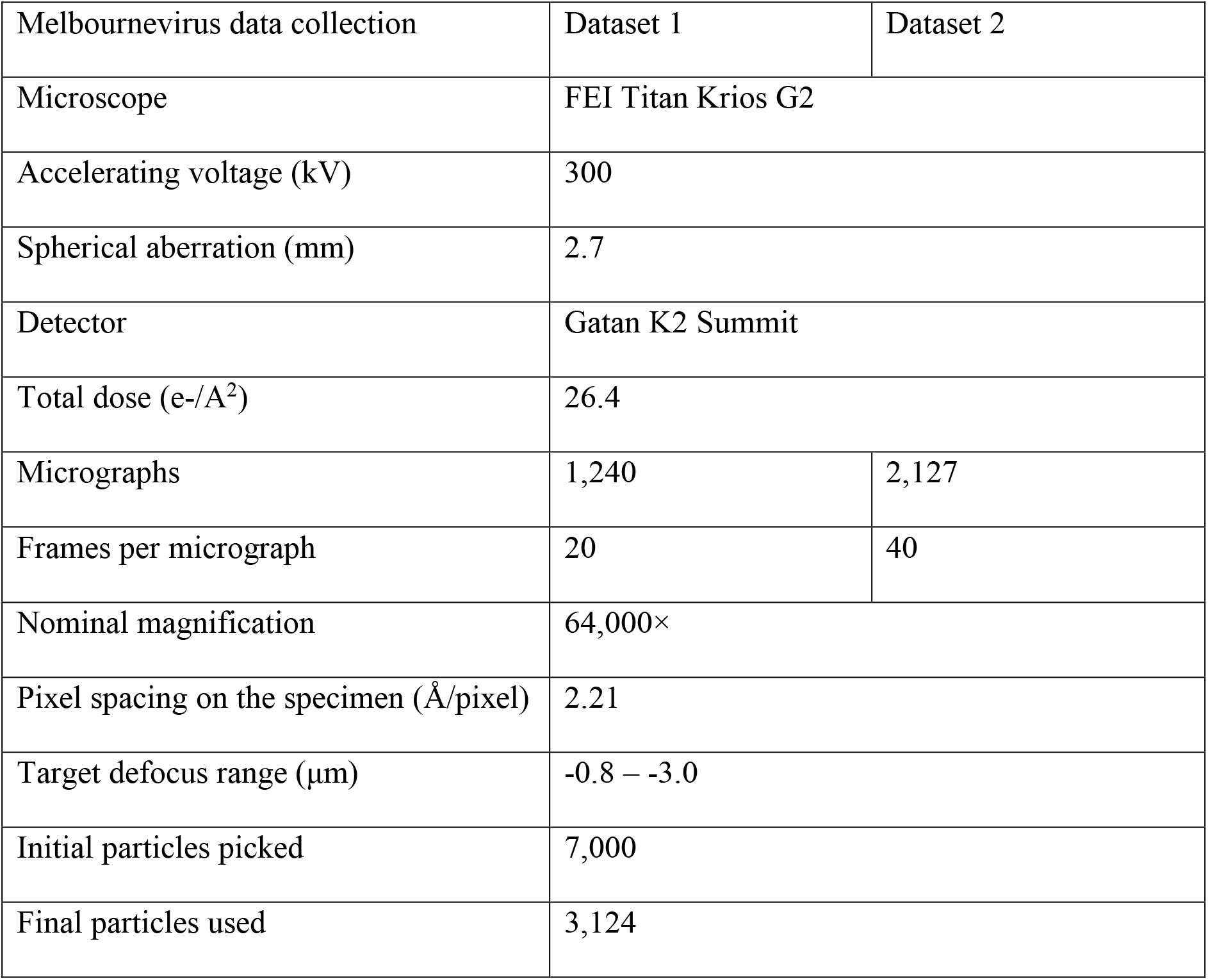
Dataset collection details

| Melbournevirus data collection | Dataset 1 | Dataset 2 |
| --- | --- | --- |
| Microscope | FEI Titan Krios G2 |  |
| Accelerating voltage (kV) | 300 |  |
| Spherical aberration (mm) | 2.7 |  |
| Detector | Gatan K2 Summit |  |
| Total dose (e-/Å<sup>2</sup>) | 26.4 |  |
| Micrographs | 1,240 | 2,127 |
| Frames per micrograph | 20 | 40 |
| Nominal magnification | 64,000× |  |
| Pixel spacing on the specimen (Å/pixel) | 2.21 |  |
| Target defocus range (μm) | -0.8 – -3.0 |  |
| Initial particles picked | 7,000 |  |
| Final particles used | 3,124 |  |

**Figure 1.**
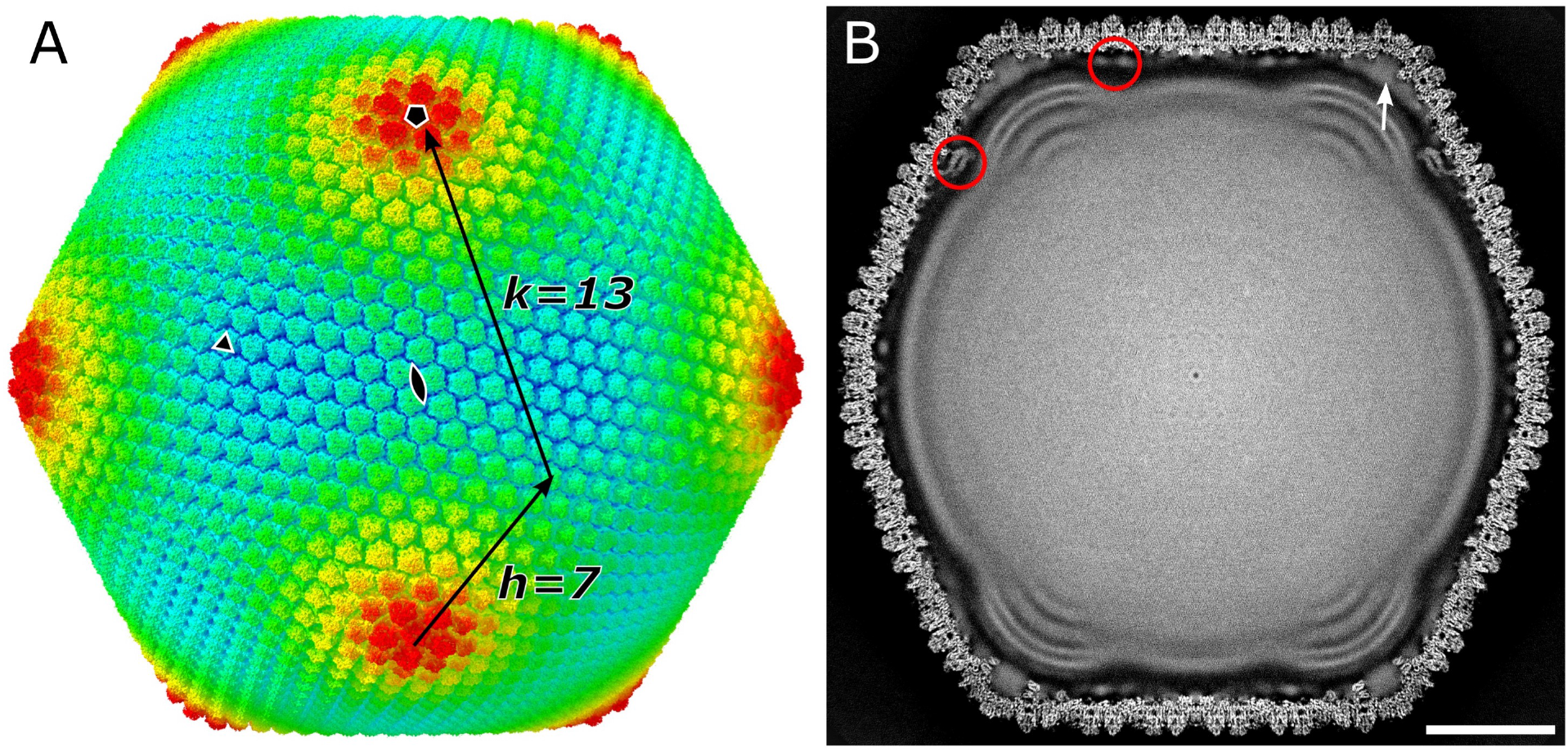
SPA 3D reconstruction of Melbournevirus at 4.9 Å. A) External view of Melbournevirus, coloured by radius with the following parameters: blue, 1,050 Å, turquoise, 1,100 Å; green, 1,150 Å; yellow, 1,200 Å; red, 1250 Å, and symmetry axes are marked by a pentagon (five-fold), triangle (three-fold) and double teardrop (two-fold). Index of h and k for triangulation number is indicated. B) Central slice of Melbournevirus. Red circles indicate the scaffold proteins between the capsid layer and inner membrane. A white arrow indicates a low resolution “plug” of density which fills a space between the pentasymmetron and the inner membrane extrusion but does not directly contact the inner membrane. Scale bar equals 50 nm.

**Figure 2.**
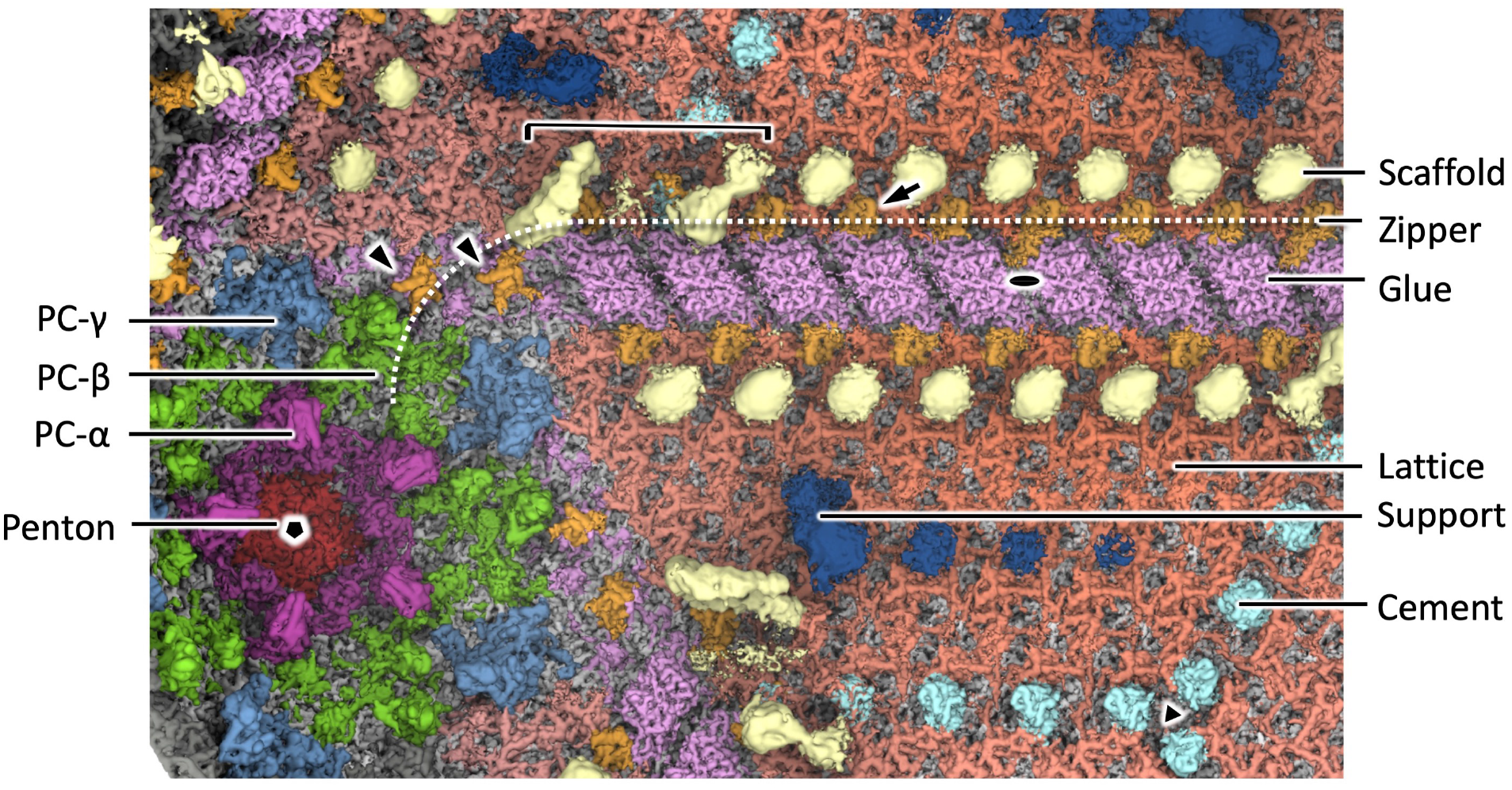
Segmentation of Melbournevirus, focussed on the asymmetric unit encompassing the five, three and twofold axes, and viewed from the inside of the capsid. The map was filtered by local resolution to permit visualisation of the flexible scaffold and support proteins, which are otherwise weak due to uniform b-factor sharpening of the map. Support, Scaffold, Zipper, Glue, Cement, Lattice, PC-α, PC-β, PC-γ, and Penton indicate the names of each mCPs mentioned in those of tokyovirus (Chihara et al., 2021). Fivefold, threefold, and twofold axes indicated by a black pentagon, triangle and double teardrop respectively. White arrow and arrowheads indicate the different orientations of the zipper protein. Bracket shows the “horseshoe” shaped head of the scaffold proteins.

With this Melbournevirus dataset, we approached the resolutions at which *de novo* model building can be attempted. Using the “block based” reconstruction strategy (Zhu et al., 2018) permitted further improvements in resolution (Fig. S2). Blocks were reconstructed with no symmetry applied (C1), with a soft mask focussed on the capsid. Carrying out the block reconstructions without defocus refinement barely improved resolution above that of the Ewald-sphere corrected whole virus reconstruction (the global FSC for the fivefold block estimated 4.8Å) but a single pass of defocus refinement resulted in the reconstructions reaching the maximum resolution of 4.42 Å (Fig. S2-S5, Table 2).

**Table 2.**
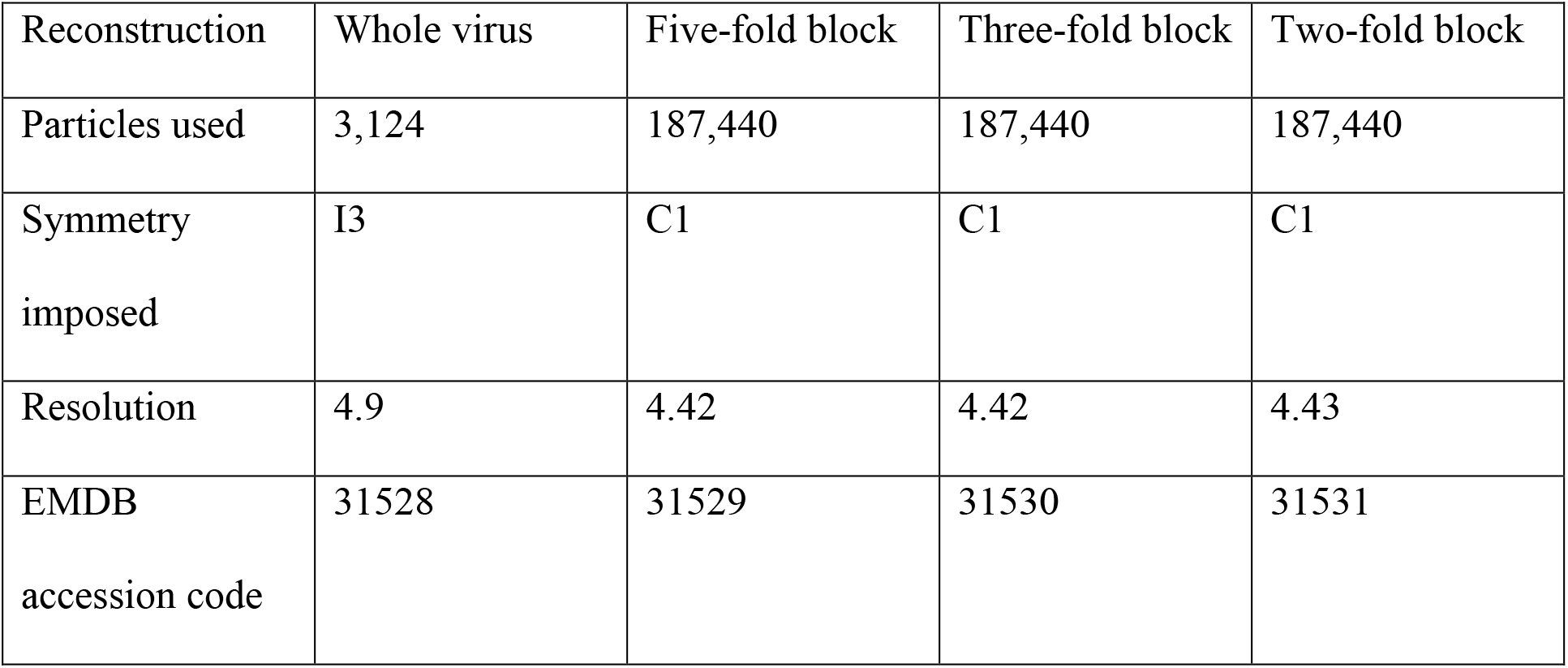
Reconstruction details

| Reconstruction | Whole virus | Five-fold block | Three-fold block | Two-fold block |
| --- | --- | --- | --- | --- |
| Particles used | 3,124 | 187,440 | 187,440 | 187,440 |
| Symmetry imposed | I3 | C1 | C1 | C1 |
| Resolution | 4.9 | 4.42 | 4.42 | 4.43 |
| EMDB accession code | 31528 | 31529 | 31530 | 31531 |

Figure 2 shows the capsid viewed from the inside, filtered by local resolution, and focussed on the asymmetric unit encompassing a two, three and fivefold symmetry axes, describing the details of the protein network formed with mCPs. Each component is labelled according to the nomenclature proposed in the segmentation of tokyovirus by Chihara *et al*. (Chihara et al., 2021) and closely follows the motifs previously described. These are individually detailed as follows.

### The molecular structure of the individual proteins

While the genome of Melbournevirus has been sequenced, most of the capsid proteins are unannotated with a few hypothetical assignments (Okamoto et al., 2018). As a result, we built a model of the MCP, and carried out chain tracing and fitting of poly-alanine chains for some other regions of density which could be segmented and followed without significant ambiguity.

#### The major capsid protein

The amino acid sequence of the Melbournevirus MCP was aligned against those of tokyovirus, PBCV-1 and other NCLDVs, where it scores highly against tokyovirus, and within acceptable parameters against Iridovirus and PBCV-1 (Fig. S6). The MCP was predicted to possess the same “double jelly roll” motif as other NCLDV MCPs (Fig. S7). This is confirmed by the striated density on the MCP trimer extracted from the centre of the threefold axis (Fig. 3A), which show a β-sheet structure. A homology model of the Melbournevirus MCP was built by using the PBCV-1 MCP (PDB ID: 5TIQ) as a structural template, and flexible-fitted to the density, clearly showing the molecular model of the MCP trimer (Fig. 3B). As previously reported for PBCV-1 (Fang et al. 2019) and ASFV (Wang et al. 2019), the MCP trimer is formed with interactions between the FG1 loops which extends inside the trimer with the two short α-helices (dotted red circle in Fig. 3D), and stabilized with the N-terminal domain (NTD) extended from the adjacent monomer which functions like an anchor (arrow in Fig. 3D). Furthermore, in the Melbournevirus MCP trimer, the elongated HI1 loop interacts with the adjacent MCP monomer (arrow heads in Fig. 3D-E, Fig. S7A), and forms a unique cup structure with the relatively long loops of DE1 and FG1 on the top of the MCP trimer (Fig. 3E). The cup structure serves to accommodate a similarly uniform and symmetric “cap” region to tokyovirus (Chihara et al., 2021) which cannot be attributed to the MCP polypeptide (Fig. 3B, C). Previously, Okamoto et al. reported that an uncharacterised protein, MEL_236, appeared to have the same abundance as the MCP protein (Okamoto et al., 2018). At 16kDa, MEL_236 is approximately the correct size (given a margin of error for SDS-PAGE size determination) to correspond with the PAS stain-sensitive 14 kDa protein of tokyovirus previously reported (Chihara et al., 2021). Genomic studies of Noumeavirus reported the orthologue of this, NMV_189, as the most abundant protein (Fabre et al., 2017). Unfortunately, in these maps the cap region is not clear enough to build a model from this sequence (arrowheads in Fig. S3D and S4D).

**Figure 3.**
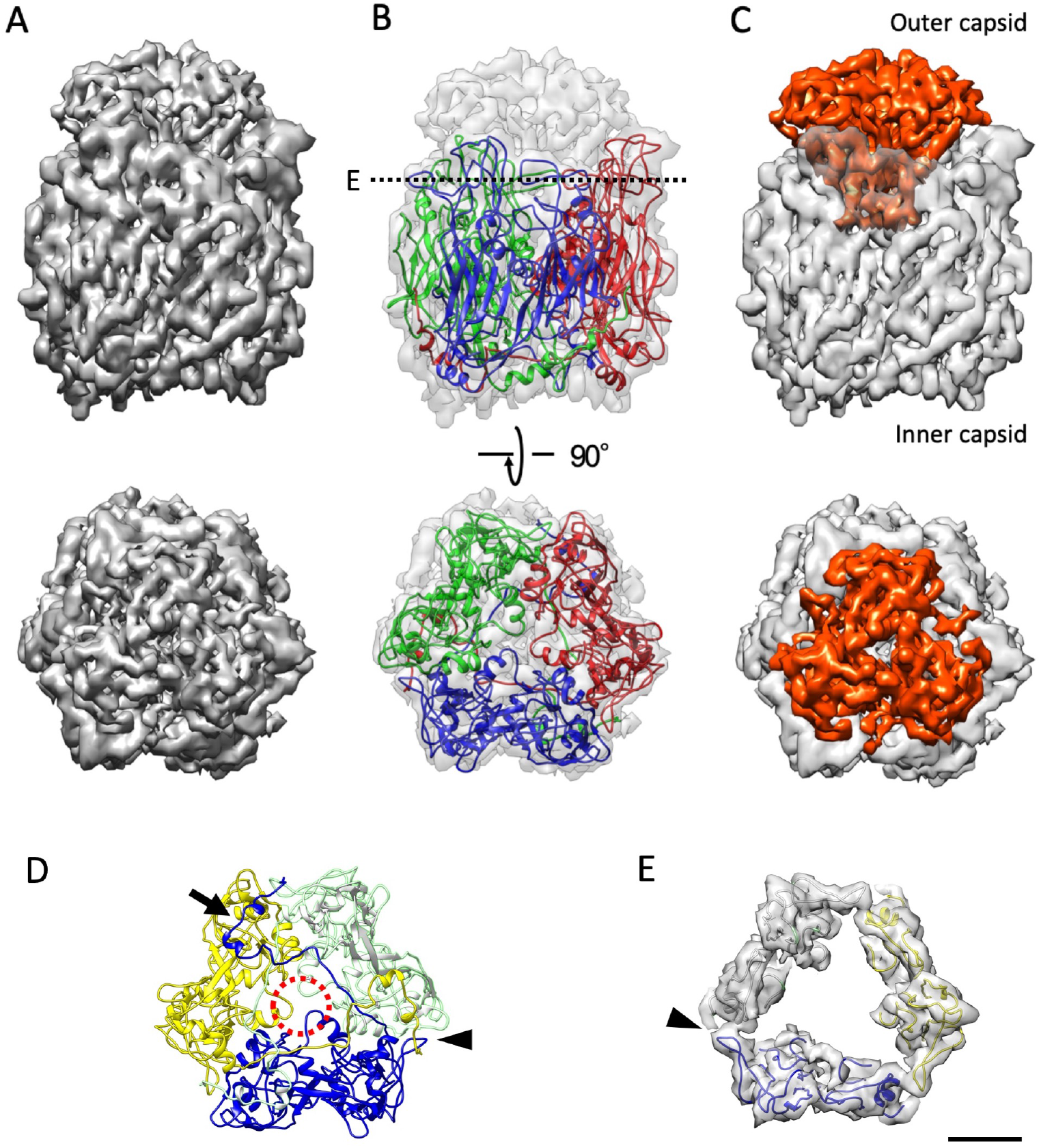
Melbournevirus MCP, with the homology model fitted. A) An MCP trimer extracted from the centre of the three-fold axis. B) Transparent density with the homology model fitted. C) The MCP with the “cap” region, which the annotated MCP sequence used for building the MCP does not fill highlighted. Top and bottom panels show the side and top view of the MCP. D) The FG1 loops (dotted red circle), N-terminal domain (arrow), and HI1 loop (arrowhead) stabilize the MCP trimer. The trimer models are viewed from the inner capsid side. E) The HI1 loop interacts with the adjacent HI2 loop and forms a cup structure with the extended DE1 and FG1 loops to accommodate the cap density (red in panel C). Scale bar equals 2 nm.

#### The penton base protein

As reported in other giant virus capsids (Chihara et al. 2021, Fang et al. 2019, Wang et al. 2019, Born et al., 2018), the five-fold axis of the Melbournevirus capsid is filled with a penton base protein pentamer. We performed a rigid body fit of the penton base protein from PBCV-1 (Fang et al., 2019) to this density, but the quality of the fit was marginal. Therefore, the penton base pentamer was extracted from the fivefold block reconstruction (Fig. S3) using UCSF Chimera (Pettersen et al., 2004) and SEGGER (Pintilie and Chiu, 2012), and DeepTracer (Pfab et al., 2021) used with this extracted density together with a short polyalanine sequence to trace the backbone (Fig. 4). The penton protein exhibited a single jelly roll (SJR) motif similar to those of other giant viruses (Fig. S8). The pentamer is formed by two interactions, external and internal; the external is the loop extended inward of the pentamer in the outer capsid region (Fig. 4C), and the internal is the interactions between the N-terminals (black asterisks in Fig. 4D) and between the N-terminal and C-terminal (red asterisks in Fig. 4D). The entire structure of the penton base protein is similar to that of PBCV-1, which does not have the large insertion domain found in CroV-dependent mavirus (Fig. S8). In the cryo-EM map, some inner capsid densities also exist under the penton base protein models (arrow in Fig. 4). Another component may exist in this region and connect the penton to the inner membrane extrusion.

**Figure 4.**
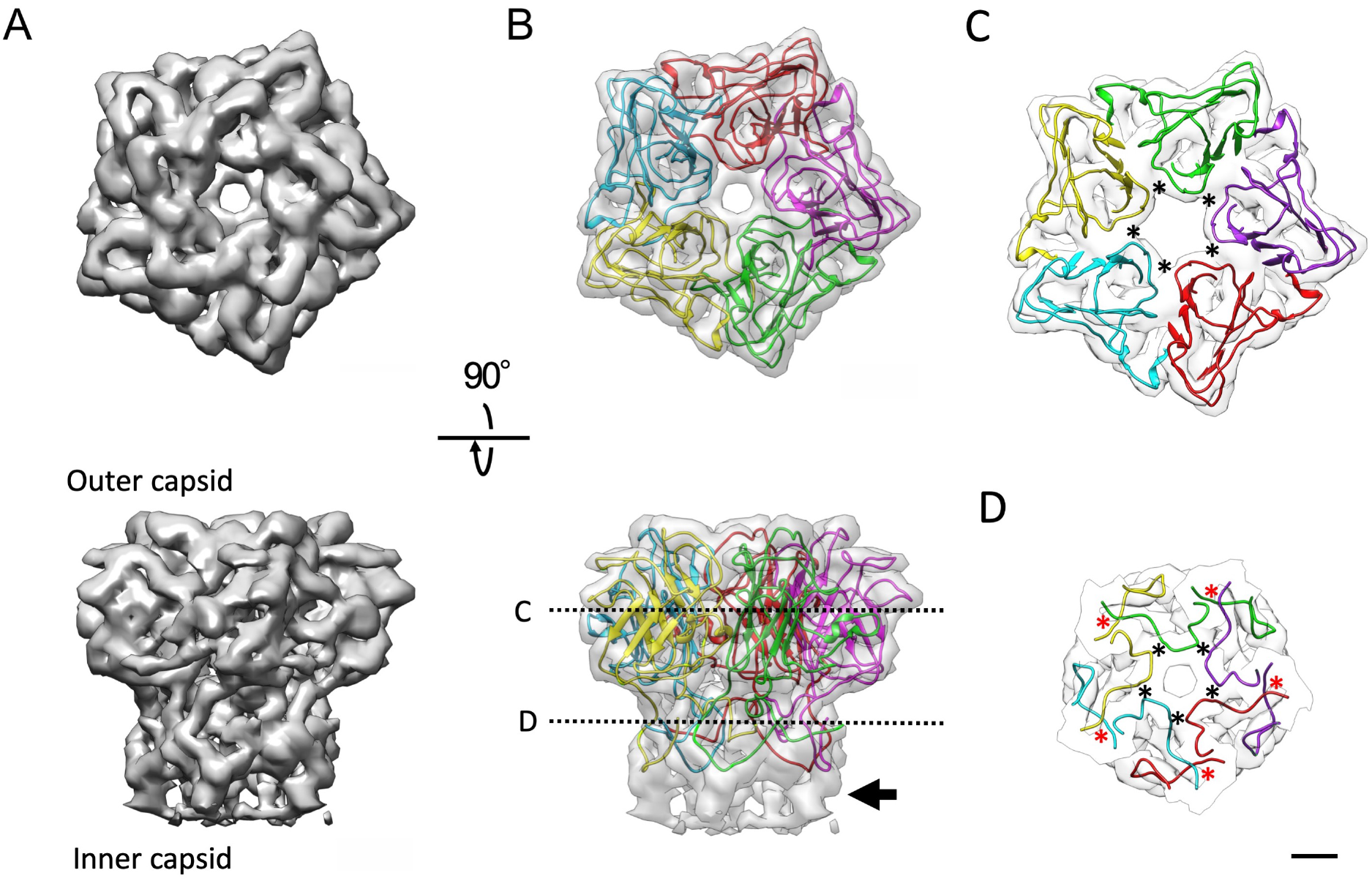
A pentamer of penton base proteins of Melbournevirus. A) The extracted penton density. B) With a *de novo* poly-alanine chain model built into the individual symmetrical densities. Top and bottom panels are viewed from the top and the side of the pentamer, respectively. Arrow indicates the density does not fill the polyalanine model. C) The loops from the individual penton base proteins externally stabilize the pentamer. The slice position is indicated in panel B. The interaction points are labeled with asterisks. D) The interactions between N- and C-terminals internally stabilize the pentamer. The slice position is indicated in panel B. The interaction points between N-terminals and between N-terminal and C-terminal are labeled with black and red asterisks, respectively. Scale bar equals 1 nm.

The five-fold axis of Melbournevirus capsid with the characteristic extrusion of the inner membrane shows a low resolution “plug” of density which fills the area underneath the penton, completely enveloping the pentasymmetron component proteins (Fig. 1B), but this plug does not directly contact the inner membrane extrusion.

#### The pentasymmetron components

As the minor capsid proteins underneath the pentasymmetron, three pentasymmetron components (PC-α, PC-β, PC-γ) have been identified in tokyovirus (Chihara et al. 2021). Similar components were also identified in Melbournevirus (Fig. 2). With the improved resolution of the Melbournevirus block-based reconstruction, we can clearly see one of the pentasymmetron component proteins (named as PC-α (Chihara et al., 2021)) revealing two quadruple bundles (each reminiscent of a ferritin monomer (Kayama et al., 2021)) of α-helices, stacked one on top of the other (Fig. 2, Fig. S9). A polyalanine model was built *de novo* and fitted into the density. However, it is less clear whether this is a single protein or two independent ones, as the extremity of this region is relatively unsupported and as a result is ∼6Å (Fig. S3). Our current hypothesis is that they are two independent bundles, where the polypeptide chain continues into the capsid framework and are located surrounding the inner membrane extrusion (Fig. S8, Fig. 1B).

#### Other minor capsid proteins

Like tokyovirus, the mCPs of Melbournevirus form an intricate lattice network (Fig. 2). The two viruses, extremely close in genetic terms, should demonstrate high structural similarity. Thus, Melbournevirus shows the same lattice array with the trapezoidal lattice consisting of the lattice protein (salmon in Fig. 2), which are linked by cement proteins (pale blue in Fig. 2) and with a glue/zipper protein arrangement (pink and orange, respectively, in Fig. 2) along the trisymmetron interface. These are internally supported by other proteins; one is the scaffold protein, and the other is the support protein (Fig. 2). The scaffold proteins (yellow in Fig. 2) extend from the “horseshoe” shaped terminus (bracket in Fig. 2) of one scaffold array near the pentasymmetron to the scaffold array along the trisymmetron interface. The support proteins (dark blue in Fig. 2) showed a large density and three additional smaller densities, running parallel to the trisymmetron interface and the cement protein array. The three additional small densities were not observed in tokyovirus capsid at 7.7 Å resolution (Chihara et al. 2021). As the scaffold and support proteins (yellow and dark blue, respectively, in Fig. 2) are underneath the lattice layer of the capsid, they are more flexible and as such resolution suffers when the map is filtered by local resolution, however without local filtering they are exceedingly difficult to see (Fig. 2). The glue proteins are located on the edge of the trisymmetron and serve to connect the adjacent trisymmetron. The bisymmetric structure looks like a dimer of the protein, but it is unclear at this resolution. The zipper protein consists of ten components (orange in Fig. 2), eight of which are located along the trisymmetron interface and bind to each of the two-fold symmetric glue proteins (pink in Fig. 2). The remaining terminal two zipper proteins are located between the glue proteins and the pentasymmetron components (arrowhead in Fig. 2), and their orientation is significantly different from the other zipper proteins (arrow in Fig. 2).

### Orientation of MCP trimers on the capsid surface

The pentasymmetron assembly of Melbournevirus shares the MCP orientation (“golf club”) motif displayed in CroV (Xiao et al. 2017) and PBCV-1 (Xian et al. 2020), in addition to the trisymmetron (Fig. 5). In the motif, the corner, P_nd_ (n=1,2,3,4,5), of the MCP trimer in the asymmetric unit shows a 60° rotation to match that of the adjacent asymmetric unit. This is curious as Xian *et al*. hypothesise this orientational variance in the pentasymmetron asymmetric unit to be caused by the tape measure protein (Xian et al. 2020), which Melbournevirus does not exhibit, but instead appears to use the scaffold array (Fig. 2) and some other mCPs which were originally reported by Chihara *et al*. (2021). Examining underneath this “golf club” motif positions PC-α and PC-β at the interfaces of the individual motifs, with PC-γ at the interface of the motif and the neighbouring trisymmetron (white circles in Fig. 5). Looking at the minor capsid protein network in the Melbournevirus capsid, the function of the tape measure proteins is possibly replaced by the coordinated use of the PC-β, zippers, and scaffold proteins (dotted curve in Fig. 2).

**Figure 5.**
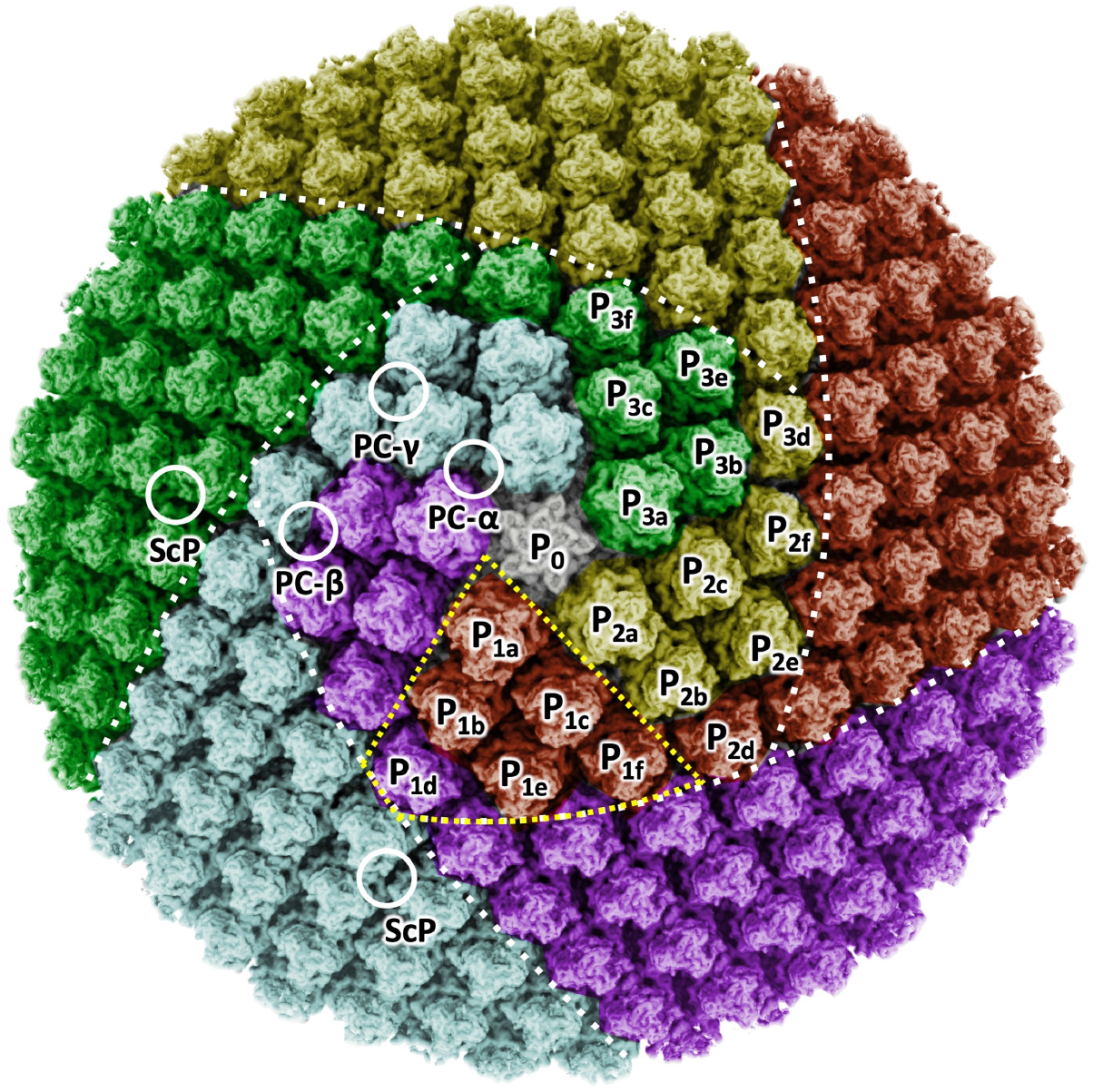
One of the pentasymmetron blocks, with MCPs coloured by orientation. The MCPs of Melbournevirus show the same orientation pattern as CroV and PVCB-1 (Xiao et al, 2017; Xian *et al*. 2020). The while dotted lines show the interfaces of trisymmetron and pentasymmetron. The yellow dotted triangle shows an asymmetric unit of the pentasymmetron. The white circles indicate the locations internal of the MCP layer where the pentasymmetron components are located (PC-α, PC-β and PC-γ respectively) and the “horseshoe” head of the scaffold protein (ScP). Three asymmetric units of the pentasymmetron are labelled P_1a-f_, P_2a-f_ and P_3a-f_, showing that one of the MCPs in the asymmetric unit (P_1-3d_) is rotated 60° to match that of the adjacent asymmetric unit.

## Discussion

Achieving close to the Nyquist limit of this dataset when reconstructing the complete viral particle with Ewald sphere correction demonstrates how vital proper analysis and correction of the various physical (electro-optical) effects can be for the study of giant viruses. The Ewald sphere reconstruction and solvent-corrected FSC curve does not fall to zero (black FSC curve, Fig. S1J). Therefore, we also show the unmasked (non-solvent corrected) Ewald sphere-corrected curve (grey curve, Fig. S1J), and the respective FSC curves without Ewald sphere correction (maroon and orange, respectively, Fig. S1J) all of which reach zero, indicating that the Ewald sphere-corrected FSC curve not reaching zero is a result of how close it is to maximum resolution. In our study, a processing method analogous to block-based reconstruction (Zhu et al., 2018) (Fig. S2-5) was used to improve the clarity of the map further. When we used 1MV cryo-HVEM for single particle analysis (SPA) of tokyovirus (Chihara et al., 2021), Ewald sphere correction did not improve the resolution. This suggests that the electron accelerated 300 kV microscope has a significant defocus gradient limitation for giant viruses above 200 nm, although symmetry expansion and block-based reconstruction helps to counter this. Further, block-based reconstruction permits subtle variation in particle orientation, which is not possible with symmetry imposition. This may also play a role in improving resolution in the block reconstructions.

This Melbournevirus cryo-EM 3D reconstruction has a comparatively low particle count of 3,124 particles when compared to the previously reported structures of PBCV-1 (∼13,000 for a 4.4 Å reconstruction (Fang et al., 2019)) and ASFV (16,266 for a 14.1 Å reconstruction (Liu et al., 2019), 63,348 for an 8.8 Å reconstruction (Wang et al., 2019)). We attribute this to both biological and computational reasons. ASFV possesses an outer membrane which will negatively impact the resolutions reported for a whole virus reconstruction due to its disordered nature, which neither PBCV-1 nor Melbournevirus possess. PBCV-1, on the other hand, achieves a higher whole virus resolution, which can be considered consistent with the increased particle count. It is also ∼25% smaller than Melbournevirus, at 190 nm, so the intra-particle defocus gradient should not be as large. Computationally, improvements in image processing software, specifically, magnification anisotropy estimation and Ewald sphere correction played a significant role in improving our Melbournevirus reconstruction (see Methods). As these computational methods advance further, we look forward to seeing further improvements for the study of giant viruses.

The Melbournevirus MCP is structurally analogous to the PBCV-1 (Fang et al., 2019), Faustovirus (Klose et al., 2016) and ASFV (Liu et al., 2019; Wang et al., 2019) MCPs, with the sequence similarity including a double jelly roll motif each composed of eight β-strands (Benson et al., 2004). The greatest level of similarity is with tokyovirus (Fig. S6), where only a few residues differ, which would suggest the MCP structure is highly preserved in *Marseilleviridae*. With the improved resolution of the Melbournevirus block reconstructions, we could better visualise the HI1 loop of Melbournevirus MCP, contacting the HI2 loop of the adjacent monomer and forming a tight trimeric complex of MCP (Fig. 3D-E, Fig. S7A). This differs from ASFV, where it is the FG1 loop which is extended (Liu et al., 2019), and from PBCV-1, where a short FG1 loop is in close proximity to the HI2 loop of the neighbouring MCP (De Castro et al., 2018). Furthermore, the HI1 loop and the other external loops of DE1 and HI2 extend externally (Fig. 3B, Fig. S7) and form a cup structure on the top of the MCP trimer to accommodate additional cap densities (orange in Fig. 3C). PBCV-1 MCP has a highly glycosylated “cap” region, while ASFV and faustovirus have a large additional β-sheet rich domain (Fig. S7B-D). Melbournevirus, however, has an ordered region, which is still lower resolution than the main body of the MCP trimer (Figs. S3D and S4D), which does not correspond to any part of the MCP model (Fig. 3B,C). This ordered structural motif would imply the presence of protein, rather than sugars, which tend to be highly mobile. However, the reduced clarity of the density compared to that of the MCP may be caused by several factors. Firstly, the cap may be in multiple conformations across the capsid surface, resulting in loss of clarity due to icosahedral averaging. Secondly, the cap may present partial occupancy of each MCP trimer across the capsid surface, also resulting in loss of clarity due to icosahedral averaging. To conclude this, further investigation is necessary to identify the cap protein and determine the stoichiometry with the MCP. As Fabre *at al*. (Fabre et al., 2017) and Okamoto *et al*. (Okamoto et al., 2018) report MEL_236 at equal or greater stoichiometry to the MCP protein, we can strongly assume that MEL_236 corresponds to the 14kDa glycosylated cap protein we previously reported (Chihara et al., 2021). Aoki et al. reported that in hokutovirus and kashiwazakivirus, which belong to lineage B of *Marseilleviridae*, the newly replicated viruses are aligned on the surface of the host cells, exhibiting a “bunch” formation consisting of normal and infected cells, similar to tupanvirus (Aoki et al. 2019). The flexible characteristics of this glycosylated cap protein on the MCP trimer may be different within the members of *Marseilleviridae*, playing a different functional role in the interface with the viral host.

With the improvement in resolution of this Melbournevirus reconstruction, we have been able to build a polyalanine model of the penton base protein. The DeepTracer (Pfab et al., 2021) chain tracing predicted a SJR fold. When comparing the DeepTracer predicted structure to that of the PBCV-1 penton base protein (Fang et al., 2019), the SJR fold aligns, but the less structured loops are of varying length. The polyalanine model fits the density well, however the loops show some dislocation from that of PBCV-1 (Fig. S8), which may explain why we have been unable to identify candidate penton protein primary sequence *via* BLAST (Altschul et al., 1990; Johnson et al., 2008) sequence homology. The CroV-dependent mavirus penton (Born et al., 2018), which we previously tested against tokyovirus, again aligns at the SJR fold but is otherwise a poor fit, even when the external insertion domain (curly bracket in Fig. S10) which is absent in PBCV-1 and Marseilleviruses, is ignored. The penton shows more density underneath the fitted polyalanine model (arrow in Fig. 4B). Weng et al. reports that the penton base pentamer hung lantern-like densities underneath the protein, connecting the penton and inner membrane (Wang et al. 2019). In the Melbournevirus penton, we do not find the lantern-like density under the penton. The low-resolution “plug” of density following the penton may have a similar function to connect the penton with the internal membrane extrusion.

There are two orientation combinations between MCP trimers, a non-rotating combination and a 60°-rotating combination (Wang et al. 2019). In the non-rotating combination, a jelly roll motif 1 (JR1) in the MCP trimer interacts with a jelly roll motif 2 (JR2) in the adjacent MCP trimer, which creates a relatively flat MCP array, like the trisymmetron surface. In contrast, in the 60°-rotating combination, a JR1 in the MCP trimer interacts with a JR1 in the adjacent MCP trimer, while a JR2 in the MCP trimer interacts with a JR2 in the adjacent MCP trimer. These combinations create a relatively angled edge, like the trisymmetron interface. In the pentasymmetron, the orientation of the MCP trimers is more complicated. Five asymmetric units of the pentasymmetron, ideally each consisting of six non-rotating combination MCP trimers, surround the central penton protein, and the interface of each asymmetric unit interacts with the 60°-rotating MCP trimer of the neighboring asymmetric unit. Finally, these MCP trimers creates a penton vertex (Fig. S11). However, Xiao et al. reported in PBCV-1 and CroV that one of six MCP trimers of the asymmetric unit, which located at the corner of the pentasymmetron, was exceptionally rotated 60° (Xiao et al. 2017, Xian et al. 2020). They called this MCP trimer array of the asymmetric unit the “golf club motif”. Furthermore, the “golf club motif” was hypothesized to be caused by specific localization of the tape measure proteins (Xian et al. 2020). In our observations, Melbournevirus showed a “golf club motif” of the MCP trimer array in the same orientation as PBCV-1 and CroV (Fig. 5), but Melbournevirus lacks a tape measure protein. However, the rotation of the single MCP trimer by 60° (P_1d_, P_2d_, P_3d_ in Fig. 5) orients it to match that of the adjacent trisymmetron and penton asymmetric unit. If we picture all the MCP trimers in a pentasymmetron asymmetric unit in the same orientation (Fig. S11), we can better visualise why the rotation of a single MCP trimer may occur. The single MCP trimer would cause a mismatch in the MCP orientational alignment of the trisymmetron interface (arrows in Fig. S11). The trisymmetron interface demonstrates a greater angle across it than the trisymmetron, which is of a gentler curvature. As such, it is likely that this orientational arrangement is a result of improving the flexibility of the capsid across the trisymmetron interface.

Without a tape measure protein, we propose that PC-β, which is present underneath the interface between the 60°-rotated P_d_ MCP trimer and the P_b_ and P_e_ MCP trimers, may play a role in determining trimer orientation. In a similar manner, PC-α may support the interface between the penton and two asymmetric units, while PC-γ supports the interface between the pentasymmetron and the adjacent trisymmetron. However, these components do not directly contact with the 60°-rotated P_d_ MCP trimer. While we have previously proposed that the scaffold protein array acts as an equivalent to the tape measure protein, the horseshoe terminus of the scaffold does not appear to be able to directly interact with any of the PC proteins, as the glue/zipper protein array is present in between the horseshoe terminus and either PC-β or PC-γ (Fig. 2). PC-β may interact with the terminal two zipper proteins, which have a rotated orientation when compared to the remaining eight zipper proteins running down the edges of the glue proteins along the trisymmetron interface, while PC-γ interacts with the glue proteins themselves along two trisymmetron interfaces. As the scaffold proteins run parallel to the glue/zipper array, some interaction may also occur there, with the three acting in concert in a similar manner to the tape measure protein in controlling capsid construction. Therefore, the combination of PC-β, zipper proteins, and scaffold proteins (dotted curve in Fig. 2) is most likely to function as the tape measure protein in Melbournevirus.

The lattice proteins show strong similarity to the motifs displayed in the lower resolution tokyovirus reconstruction. By using local resolution filtering in RELION 3.1 (Zivanov et al., 2020), we were able to avoid losing clarity in the more rigid lattice proteins, while simultaneously visualising the more flexible scaffold array (Fig. 2). Interestingly, this permitted visualisation of three additional weak densities extending from the previously reported support protein (dark blue in Fig. 2), which run parallel to the zipper/glue proteins and cement proteins on the trapezoidal lattice, but which themselves were not previously observed in tokyovirus. Missing them in tokyovirus was likely due to resolution limitations and caused by decreased signal to noise from fewer particles and lack of dose weighting for HVEM data (Chihara et al., 2021). These may act to further support the lattice array as the array extends from the pentasymmetron construct along the trisymmetron interface, in addition to supporting the characteristic internal membrane structure.

Unfortunately, while at 4.42 Å resolution we can see some secondary structure, most if not all residue sidechains are unclear. Furthermore, at many points it is extremely difficult to determine which direction of two apparently divergent paths the density should follow. As the primary sequence is unknown, and large or interacting sidechains can create a density bridge which is difficult to discriminate from backbone density, we erred on the side of caution for the time being. One of the mCPs, a repeating segment of the lattice protein (salmon in Fig. 2), has a section exposed on the internal face of the capsid (following the chain into the mCP layer created too much ambiguity in direction of the polypeptide) which fits a short section consisting of three linked α-helices (Fig. S10).

Here we have increased the resolution of a cryo-EM 3D reconstruction of a Marseillevirus to a resolution approaching that necessary to build *de novo* structural models. This has permitted us to improve our previous Marseillevirus MCP model and begin the process of building a whole-capsid model. This is currently limited by both a lack of comprehensive annotation of capsid-related proteins and reaching the maximum resolution possible with this dataset. To fully clarify the structure of Melbournevirus, and build a complete model of the *Marseilleviridae* capsid, we must fully utilize our understanding of the genetics of the *Marseilleviridae*, using copy numbers to help to identify the structural components and selected knockout genes to retrieve a phenotype, whilst simultaneously achieving resolutions that allow *ab initio* structure determination.

## Methods

### Viral propagation and purification

Melbournevirus particles were propagated and purified as described previously (Doutre et al., 2014; Okamoto et al., 2018). It is described briefly as follows. Seed Melbournevirus particles were propagated in *Acanthamoeba castellanii* cells that were cultured in PPYG medium and confluent in ten 75 cm^3^ culture flasks. The infected culture fluid was collected three days post infection and centrifuged for 10 min at 500 *g*, 4°C. The supernatant was centrifuged for 35 min at 6,500 *g*, 4°C, and the pellet was suspended in 1 mL of PBS (-). The sample was ultracentrifuged in 10-60% (w/v) continuous sucrose gradient for 90 min at 6,500 *g*, 4°C. The virus particles’ band was collected from the sucrose gradient, and centrifuged for 35 min at 6,500 *g*, 4°C. The pellet was rinsed five times in 50 mL of PBS (-) by centrifugation to remove excess sucrose. Finally, the virus particles were suspended in 100 µL PBS (-) for further cryo-EM analysis.

### Cryo-EM sample preparation and acquisition

Three µL of purified virus particles were loaded on a glow-charged lacey carbon copper grid and flash frozen in liquid ethane after 2 s blotting in 100% humidity at 4°C using a Vitrobot Mark IV (Thermo Fisher Scientific). The particles images were collected using a 300 kV Titan Krios G2 microscope (Thermo Fisher Scientific) equipped with a K2 Summit direct electron detector using counting mode and a GIF energy filter of 20 eV slit width (Gatan Inc). Two datasets were collected, dataset 1 with 20 frames per micrograph movie, and dataset 2 with 40 frames per micrograph movie. These datasets were taken at a nominal magnification of 64,000× (pixel size on specimen is 2.21 Å/pixel), at 0.8 – 3.0 µm under focus, and at 26.4 e^-^/Å^2^ total dose.

### Image processing

The image processing workflow of the whole virus particle in summarised graphically in Fig. S1. 3,367 micrograph movies were imported into RELION 3.1 (Zivanov et al., 2018; Zivanov et al., 2020) and motion correction performed using MotionCor2 (1.4.0) (Zheng et al., 2017) using whole frame motion correction. Contrast transfer function (CTF) parameters were calculated on the resulting micrographs using CTFFIND 4.1.14 (Rohou and Grigorieff, 2015), using the RELION 3.1 default parameters except for defocus step size, which was set to 100 Å. A total of 229 particles were manually picked, extracted using 4× downsampling (to 8.84 Å/pixel in 360-pixel boxes), and 2D classified into 10 classes with 0.5° angular sampling and “ignore CTFs until first peak” enabled. Three classes were used to generate three initial models using I3 symmetry with a mask diameter of 2550 Å and an initial angular sampling of 3.7°, of which the best was chosen. These three clear classes, comprising 182 particles, were used as references for RELION template autopicking of the full dataset, resulting in 7,000 particles picked. These were extracted using 4× downsampling and 2D classified into 40 classes with a mask diameter of 2650 Å, angular sampling of 1° and “ignore CTFs until first peak” enabled. 19 classes, totalling 5,047 particles were selected in clear classes and subjected to a further round of 2D classification, this time with 0.5° angular sampling and full CTF correction. This resulted in 3 clear classes, containing 3,852 particles. 3D classification into five classes was carried out with an angular sampling of 1.8° and “ignore CTFs until first peak” enabled. Three classes, totalling 3,569 particles, were selected from the 3D classification and a 3D refinement carried out. The maximum resolution possible with the downsampled particles was achieved (17.68 Å). Particles were re-extracted using 2× downsampling (4.42 Å/pixel in 720-pixel boxes) and a further 3D refinement and single round of per-particle defocus refinement again achieved the maximum resolution possible with downsampled particles (8.84 Å). Particles were re-extracted without downsampling (2.21 Å/pixel) which permitted a reconstruction to achieve 7 Å. Unfortunately, CTF refinement with 1,440-pixel boxes would fail. Particles were re-extracted with 1.5× downsampling (3.315 Å/pixel in 960-pixel boxes) and anisotropic magnification distortion estimated, followed by further defocus refinement. This resulted in a 6.65 Å reconstruction. 3D classification with alignment disabled was carried out into five classes, with the two best classes, comprising 3,124 particles, selected, resulting in a 6.45 Å reconstruction. Ewald sphere correction was used with *relion_reconstruct* to generate independent half-maps using the parameters from this final refinement, which results in a final gold standard FSC (Chen et al., 2013) global estimated resolution of 4.9 Å with a soft capsid only mask.

At this point, we moved toward a strategy akin to that of block-based reconstruction (Zhu et al., 2018), but only using the functions contained within RELION 3.1 (Zivanov et al., 2020). The image processing workflow of the block-based reconstruction is summarised in Fig. S2. First, symmetry expansion was carried out. Then three masks, individually focussed on a five-, three- and two-fold axis were generated in UCSF Chimera (Pettersen et al., 2004), before particle subtraction was used to calculate the shifts to centre the focussed region in the box. Particle subtraction was cancelled after shifts were calculated. Using these RELION-calculated shifts, the five-, three- and two-fold axes were extracted in 440-, 500- and 700-pixel boxes respectively, each containing a total of 187,440 boxes. All block reconstructions were carried out with C1 (no) symmetry imposed. The *relion_reconstruct* module was used for each extracted particle set to generate a reference for refinement and check positioning was correct. 3D refinements were carried out for each segment, achieving 4.8-5 Å with post-processing. Defocus refinement was carried out once for each, and *relion_reconstruct* used to reconstruct half-maps using the defocus-refined particle parameters. The five-fold and three-fold reconstructions achieved 4.42 Å, the maximum resolution possible with these data. The two-fold reconstruction achieved 4.43 Å, likely caused by the larger box size used. Further CTF parameter optimisation and Bayesian polishing were not carried out.

### Model building

The amino acid sequence of the MCP of Melbournevirus is annotated, along with four other candidates which may be mCPs previously reported by Okamoto *et al*. (Okamoto et al., 2018). A homology model of the MCP was generated using SWISS-MODEL (Waterhouse et al., 2018) using a structural template of that of PBCV-1 (PDBID: 5TIQ). This model was rigid body fit into a SEGGER-extracted (Pintilie and Chiu, 2012) MCP from the centre of the threefold axis and fitted into the map using ISOLDE (Croll, 2018) with restraints on predicted secondary structure. Real-space refinement was carried out in PHENIX (Adams et al., 2010) with intermediate rounds of geometry minimisation.

The PDB model of the PBCV-1 penton protein was rigid-body fit to the Melbournevirus penton density. It was deemed a poor fit, so the map blocks were segmented using SEGGER (Pintilie and Chiu, 2012) and extracted segments were processed with DeepTracer (Pfab et al., 2021) using a poly-alanine sequence for chain tracing. The best of these poly-alanine chain models was rigid-body fit to fill the penton with five identical chains. ISOLDE (Croll, 2018) was used to better fit some sections of the chain. Real-space refinement and validation were carried out in PHENIX (Adams et al., 2010). Other parts of the mCPs (PC-α and Lattice protein) were also process with the same way and the polyalanine models were built.

### Data availability

The density maps have been deposited in the EMDB with the accession codes 31528 (whole virus), 31529 (five-fold block), 31530 (three-fold block) and 31531 (two-fold block).

## Acknowledgements

We thank Marta Carroni at SciLifeLab cryo-EM facility in Stockholm for data collection. This study was funded by the following grants: MEXT KAKENHI (JP19H04845 to K.M.), the Joint Research of ExCELLS (20-004 to K.M.), Vetenskapsrådet (VR)/The Swedish Research Council (to K.O., grant no. 2018-03387), the Swedish Foundation for International Cooperation in Research and Higher Education (STINT) (to Janos Hajdu and K.O., grant no. JA2014-5721), FORMAS research grant from the Swedish Research Council for Environment, Agricultural Sciences, and Spatial Planning (to K.O., grant no. 2018-00421), the Royal Swedish Academy of Sciences (to K.O., grant no. BS2018-0053).

## Author contributions

K.O. and K.M. conceived the project. C.A. and M.S. provided Melbournevirus sample. H.K.N.R. and K.O. prepared cryo-EM sample. H.K.N.R. and K.O. collected cryo-EM data. R.N.B.S. processed cryo-EM data. R.N.B.S., and K.M. prepared figures and wrote the main manuscript text. All authors reviewed the manuscript.

## References

Adams, P.D., Afonine, P.V., Bunkoczi, G., Chen, V.B., Davis, I.W., Echols, N., Headd, J.J., Hung, L.W., Kapral, G.J., Grosse-Kunstleve, R.W., et al. (2010). PHENIX: a comprehensive Python-based system for macromolecular structure solution. Acta Crystallogr D Biol Crystallogr 66, 213–221.

Aherfi, S., Colson, P., Audoly, G., Nappez, C., Xerri, L., Valensi, A., Million, M., Lepidi, H., Costello, R., and Raoult, D. (2016). Marseillevirus in lymphoma: a giant in the lymph node. Lancet Infect Dis 16, e225–e234.

Altschul, S.F., Gish, W., Miller, W., Myers, E.W., and Lipman, D.J. (1990). Basic local alignment search tool. J Mol Biol 215, 403–410.

Aoki, K., Hagiwara, R., Akashi, M., Sasaki, K., Murata, K., Ogata, H., and Takemura, M. (2019). Fifteen Marseilleviruses Newly Isolated From Three Water Samples in Japan Reveal Local Diversity of Marseilleviridae. Front Microbiol 10, 1152.

Benson, S.D., Bamford, J.K., Bamford, D.H., and Burnett, R.M. (2004). Does common architecture reveal a viral lineage spanning all three domains of life? Mol Cell 16, 673–685.

Born, D., Reuter, L., Mersdorf, U., Mueller, M., Fischer, M.G., Meinhart, A., and Reinstein, J. (2018). Capsid protein structure, self-assembly, and processing reveal morphogenesis of the marine virophage mavirus. Proc Natl Acad Sci U S A 115, 7332–7337.

Boyer, M., Yutin, N., Pagnier, I., Barrassi, L., Fournous, G., Espinosa, L., Robert, C., Azza, S., Sun, S., Rossmann, M.G., et al. (2009). Giant Marseillevirus highlights the role of amoebae as a melting pot in emergence of chimeric microorganisms. Proc Natl Acad Sci U S A 106, 21848–21853.

Brandes, N., and Linial, M. (2019). Giant Viruses-Big Surprises. Viruses 11, 404.

Chen, S., McMullan, G., Faruqi, A.R., Murshudov, G.N., Short, J.M., Scheres, S.H., and Henderson, R. (2013). High-resolution noise substitution to measure overfitting and validate resolution in 3D structure determination by single particle electron cryomicroscopy. Ultramicroscopy 135, 24–35.

Chihara, A., Burton-Smith, R.N., Kajimura, N., Mitsuoka, K., Okamoto, K., Song, C., and Murata, K. (2021). A novel capsid protein network allows the characteristic inner membrane structure of Marseilleviridae giant viruses. bioRxiv, 2021.2002.2003.428533.

Colson, P., Pagnier, I., Yoosuf, N., Fournous, G., La Scola, B., and Raoult, D. (2013). “Marseilleviridae”, a new family of giant viruses infecting amoebae. Archives of Virology 158, 915–920.

Croll, T.I. (2018). ISOLDE: a physically realistic environment for model building into low-resolution electron-density maps. Acta Crystallogr D Struct Biol 74, 519–530.

De Castro, C., Klose, T., Speciale, I., Lanzetta, R., Molinaro, A., Van Etten, J.L., and Rossmann, M.G. (2018). Structure of the chlorovirus PBCV-1 major capsid glycoprotein determined by combining crystallographic and carbohydrate molecular modeling approaches. Proc Natl Acad Sci U S A 115, E44–E52.

Doutre, G., Philippe, N., Abergel, C., and Claverie, J.M. (2014). Genome analysis of the first Marseilleviridae representative from Australia indicates that most of its genes contribute to virus fitness. J Virol 88, 14340–14349.

Fabre, E., Jeudy, S., Santini, S., Legendre, M., Trauchessec, M., Couté, Y., Claverie, J.-M., and Abergel, C. (2017). Noumeavirus replication relies on a transient remote control of the host nucleus. Nat Commun 8, 15087.

Fang, Q., Zhu, D., Agarkova, I., Adhikari, J., Klose, T., Liu, Y., Chen, Z., Sun, Y., Gross, M.L., Van Etten, J.L., et al. (2019). Near-atomic structure of a giant virus. Nat Commun 10, 388.

Fernandez-Leiro, R., and Scheres, S.H.W. (2017). A pipeline approach to single-particle processing in RELION. Acta Crystallogr D Struct Biol 73, 496–502.

Iyer, L.M., Aravind, L., and Koonin, E.V. (2001). Common origin of four diverse families of large eukaryotic DNA viruses. J Virol 75, 11720–11734.

Johnson, M., Zaretskaya, I., Raytselis, Y., Merezhuk, Y., McGinnis, S., and Madden, T.L. (2008). NCBI BLAST: a better web interface. Nucleic Acids Res 36, W5–9.

Kayama, Y., Burton-Smith, R.N., Song, C., Terahara, N., Kato, T., and Murata, K. (2021). Below 3 A structure of apoferritin using a multipurpose TEM with a side entry cryoholder. Sci Rep 11, 8395.

Klose, T., Reteno, D.G., Benamar, S., Hollerbach, A., Colson, P., La Scola, B., and Rossmann, M.G. (2016). Structure of faustovirus, a large dsDNA virus. Proc Natl Acad Sci U S A 113, 6206–6211.

Kutikhin, A.G., Yuzhalin, A.E., and Brusina, E.B. (2014). Mimiviridae, Marseilleviridae, and virophages as emerging human pathogens causing healthcare-associated infections. GMS hygiene and infection control 9, Doc16.

Liu, S., Luo, Y., Wang, Y., Li, S., Zhao, Z., Bi, Y., Sun, J., Peng, R., Song, H., Zhu, D., et al. (2019). Cryo-EM Structure of the African Swine Fever Virus. Cell Host Microbe 26, 836–843 e833.

Okamoto, K., Miyazaki, N., Reddy, H.K.N., Hantke, M.F., Maia, F., Larsson, D.S.D., Abergel, C., Claverie, J.M., Hajdu, J., Murata, K., et al. (2018). Cryo-EM structure of a Marseilleviridae virus particle reveals a large internal microassembly. Virology 516, 239–245.

Pettersen, E.F., Goddard, T.D., Huang, C.C., Couch, G.S., Greenblatt, D.M., Meng, E.C., and Ferrin, T.E. (2004). UCSF Chimera--a visualization system for exploratory research and analysis. J Comput Chem 25, 1605–1612.

Pfab, J., Phan, N.M., and Si, D. (2021). DeepTracer for fast de novo cryo-EM protein structure modeling and special studies on CoV-related complexes. Proc Natl Acad Sci U S A 118.

Pintilie, G., and Chiu, W. (2012). Comparison of Segger and other methods for segmentation and rigid-body docking of molecular components in cryo-EM density maps. Biopolymers 97, 742–760.

Rohou, A., and Grigorieff, N. (2015). CTFFIND4: Fast and accurate defocus estimation from electron micrographs. J Struct Biol 192, 216–221.

Scheres, S.H.W. (2012). RELION: Implementation of a Bayesian approach to cryo-EM structure determination. J Struct Biol 180, 519–530.

Subramaniam, K., Behringer, D.C., Bojko, J., Yutin, N., Clark, A.S., Bateman, K.S., van Aerle, R., Bass, D., Kerr, R.C., Koonin, E.V., et al. (2020). A New Family of DNA Viruses Causing Disease in Crustaceans from Diverse Aquatic Biomes. mBio 11, e02938.

Wang, N., Zhao, D., Wang, J., Zhang, Y., Wang, M., Gao, Y., Li, F., Wang, J., Bu, Z., Rao, Z., et al. (2019). Architecture of African swine fever virus and implications for viral assembly. Science 366, 640–644.

Waterhouse, A., Bertoni, M., Bienert, S., Studer, G., Tauriello, G., Gumienny, R., Heer, F.T., de Beer, T.A.P., Rempfer, C., Bordoli, L., et al. (2018). SWISS-MODEL: homology modelling of protein structures and complexes. Nucleic Acids Res 46, W296–W303.

Xian, Y., Avila, R., Pant, A., Yang, Z., and Xiao, C. (2020). The Role of Tape Measure Protein in Nucleocytoplasmic Large DNA Virus Capsid Assembly. Viral Immunol 34, 41–48.

Xiao, C., Fischer, M.G., Bolotaulo, D.M., Ulloa-Rondeau, N., Avila, G.A., and Suttle, C.A. (2017). Cryo-EM reconstruction of the Cafeteria roenbergensis virus capsid suggests novel assembly pathway for giant viruses. Sci. Rep. 7, 5484.

Zheng, S.Q., Palovcak, E., Armache, J.P., Verba, K.A., Cheng, Y., and Agard, D.A. (2017). MotionCor2: anisotropic correction of beam-induced motion for improved cryo-electron microscopy. Nat Methods 14, 331–332.

Zhu, D., Wang, X., Fang, Q., Van Etten, J.L., Rossmann, M.G., Rao, Z., and Zhang, X. (2018). Pushing the resolution limit by correcting the Ewald sphere effect in single-particle Cryo-EM reconstructions. Nat Commun 9, 1552.

Zivanov, J., Nakane, T., Forsberg, B.O., Kimanius, D., Hagen, W.J., Lindahl, E., and Scheres, S.H. (2018). New tools for automated high-resolution cryo-EM structure determination in RELION-3. Elife 7, e42166.

Zivanov, J., Nakane, T., and Scheres, S.H.W. (2020). Estimation of high-order aberrations and anisotropic magnification from cryo-EM data sets in RELION-3.1. IUCrJ 7, 253–267.

