## Supplementary information for "The 4.4 Å structure of the giant Melbournevirus virion belonging to the *Marseilleviridae* family"

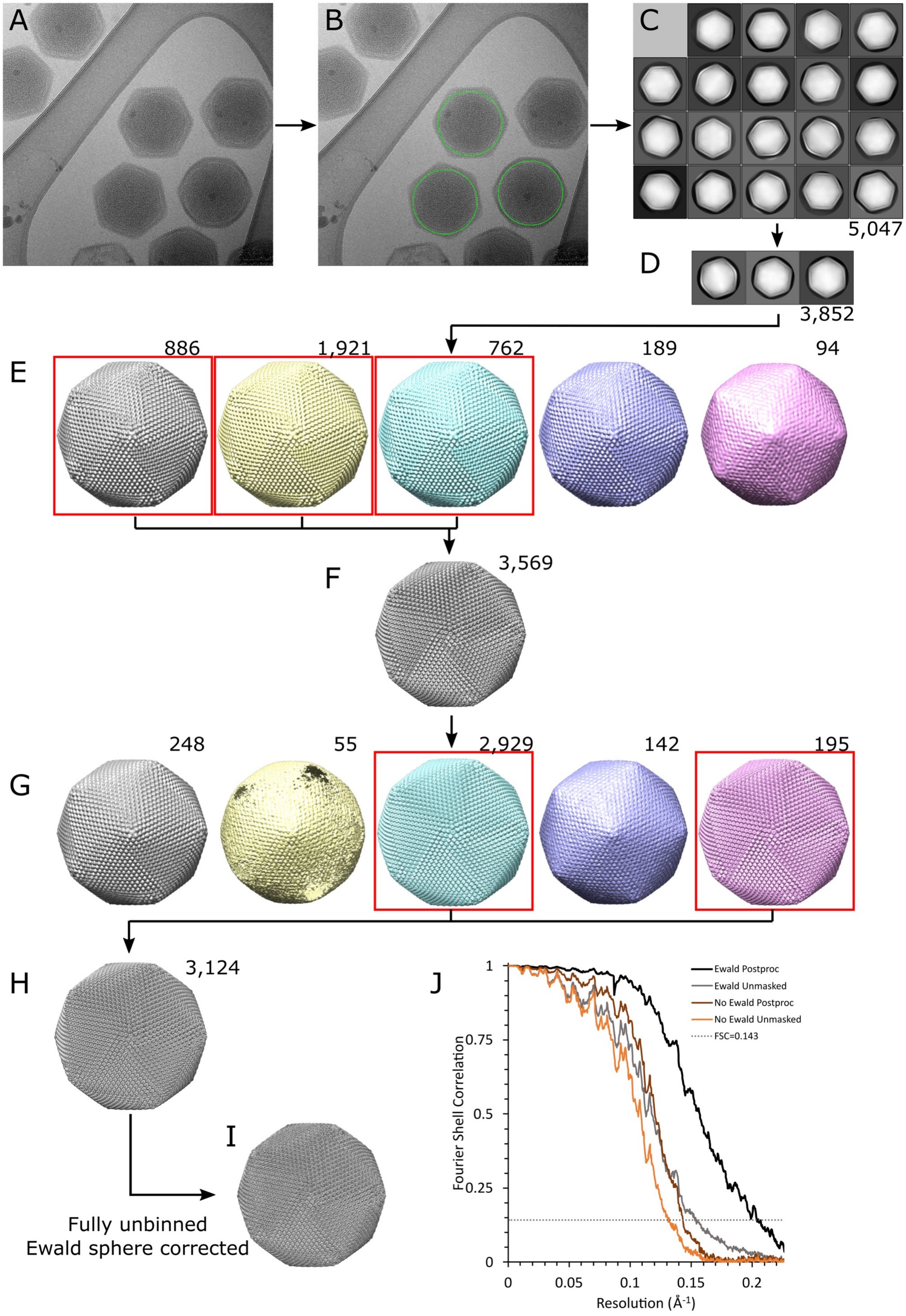

A
B
C
5,047
D
3,852
E
886
1,921
762
189
94
F
3,569
G
248
55
2,929
142
195
H
3,124
I
Fully unbinned
Ewald sphere corrected
J
Ewald Postproc
Ewald Unmasked
No Ewald Postproc
No Ewald Unmasked
FSC=0.143
Fourier Shell Correlation
Resolution (Å^-1)


24

**Figure S1.** Processing flowchart for the Melbournevirus whole-virus reconstruction. All processing was carried out in RELION (Fernandez-Leiro and Scheres, 2017; Scheres, 2012; Zivanov et al., 2018; Zivanov et al., 2020). See methods for a full description of processing. A) Representative micrograph. B) Representative micrograph with selected particles indicated with a green box. A total of 7,000 particles were autopicked. C) Selected 2D classes from the first round of 2D classification. The images are extracted with 4× downsampling. D) Selected 2D classes from the second round of 2D classification. E) The initial 3D classification into five classes, with selected classes boxed in red. F) The first 3D reconstruction from the selected particles. G) A final 3D classification, selected classes boxed in red. H) 3D refinement before fully unbinning particles. I) Ewald sphere corrected final reconstruction. J) Global gold-standard Fourier shell correlation curves of the final reconstruction masked and unmasked, with and without Ewald sphere correction. Numbers by (C, D) indicate total selected particles. Numbers by 3D models (E-H) indicate number of particles within that class.

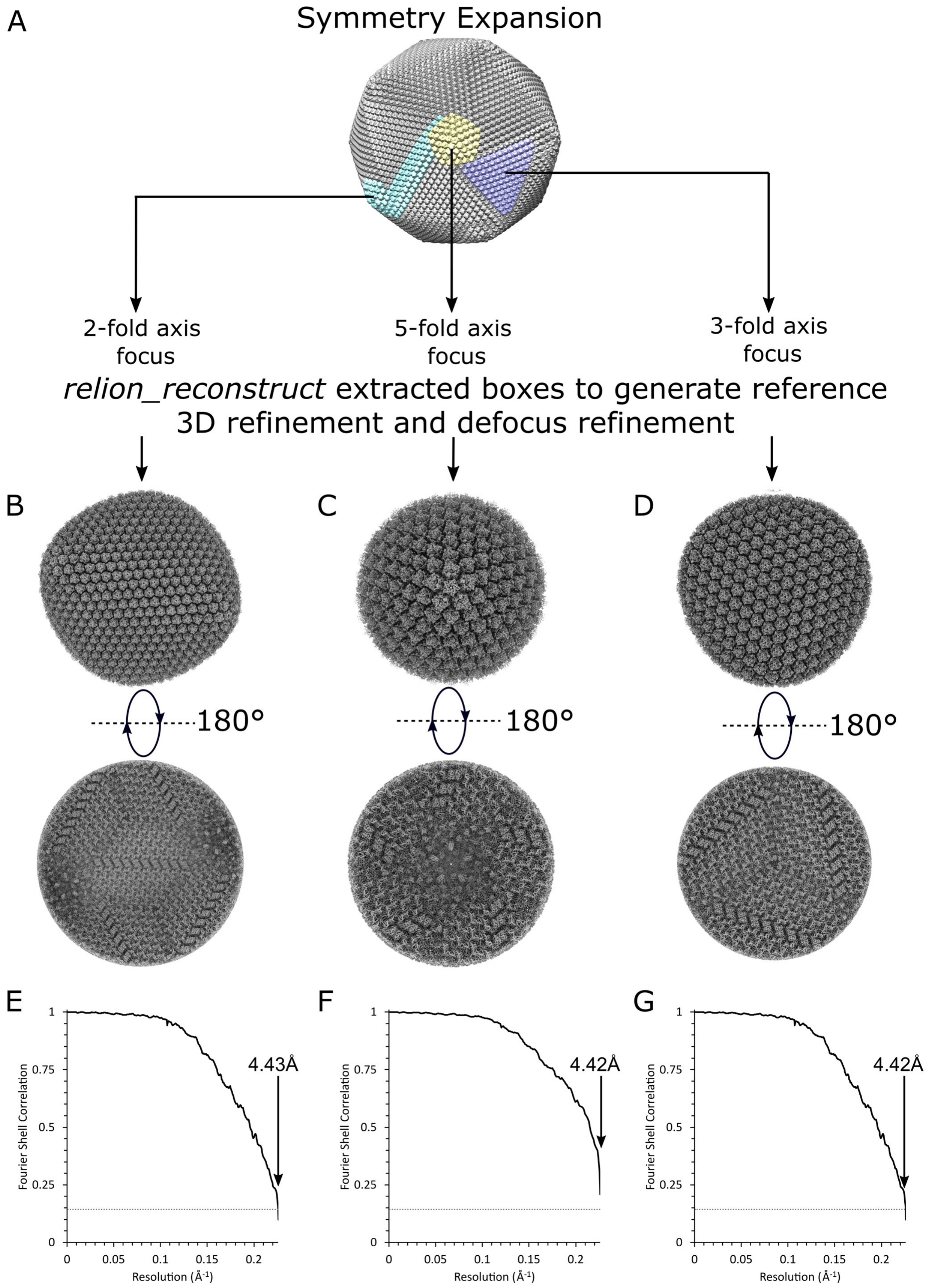
A
Symmetry Expansion
2-fold axis focus
5-fold axis focus
3-fold axis focus
relion_reconstruct extracted boxes to generate reference
3D refinement and defocus refinement
B
C
D
180°
180°
180°
E
F
G
Fourier Shell Correlation
Resolution (Å-1)
4.43Å
4.42Å
4.42Å


39

**Figure S2.** Processing flowchart of the Melbournevirus block-based reconstructions. See methods for a full description of processing. A) The 4.9 Å whole-virus reconstruction, with the five-fold, three-fold and two-fold axes masked and coloured in yellow, purple, and cyan, respectively. Symmetry expansion was carried out resulting in 187,440 particles for each block, and the 3D refinements of the blocks were carried out. A single pass of defocus refinement was used before a final reconstruction. B) two-fold axis 3D reconstruction, external and internal views. C) Five-fold axis 3D reconstruction, external and internal views. D) Three-fold axis 3D reconstruction, external and internal views. E, F, G) Gold-standard Fourier shell correlation curves for the two-fold axis block, the five-fold axis block, and the three-fold axis block, respectively. All defocus-refined block reconstructions achieve Nyquist frequency.

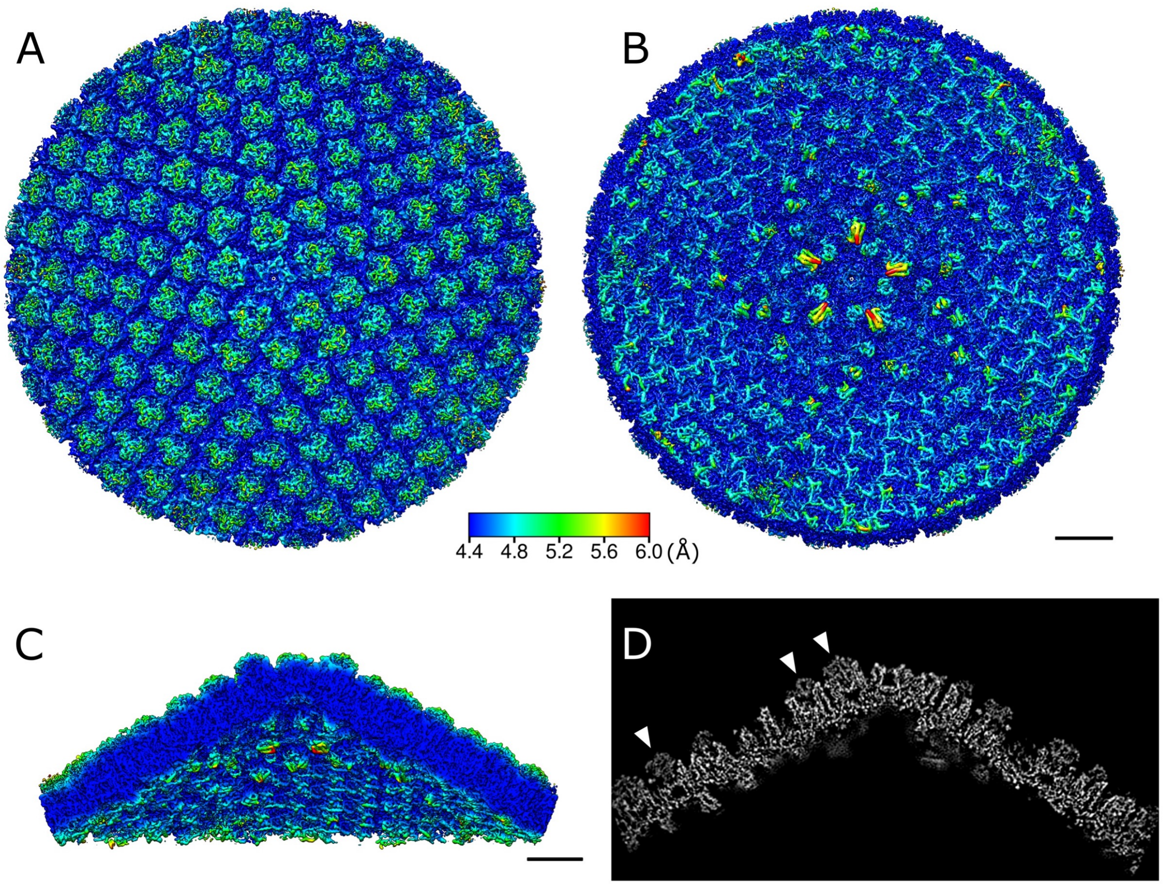


**Figure S3.** Block-based reconstruction of a five-fold axis point. A) External view. B) Internal view. C) Slice view perpendicular to (A) and (B). The map is colour coded according to the local resolution of the area. The colour code is shown in the figure. D) Slice view of density. White arrowheads indicate a centrally sliced MCP, showing the “cup” in the MCP which is filled with a cap density (arrowheads). Scale bars equal 10 nm.

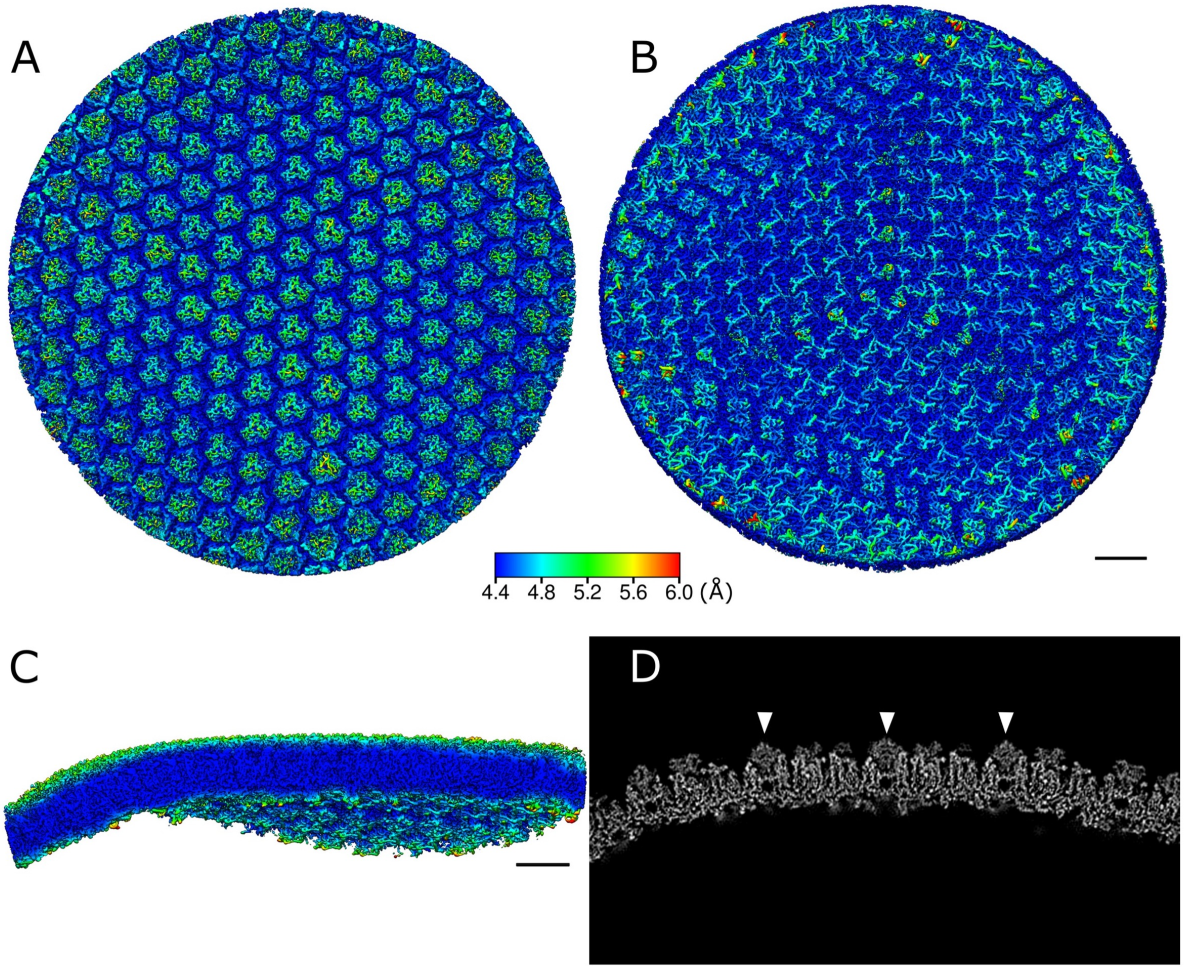


**Figure S4.** Block-based reconstruction of a three-fold axis point. A) External view. B) Internal view. C) Slice view perpendicular to (A) and (B). The map is colour coded according to the local resolution of the area. The colour code is shown in the figure. D) Slice view of density map. White arrowheads indicate a centrally sliced MCP, showing strong density for main MCP body and weaker density for the cap region (arrowheads), which fills the “cup” of the MCP. Scale bars equal 10 nm.

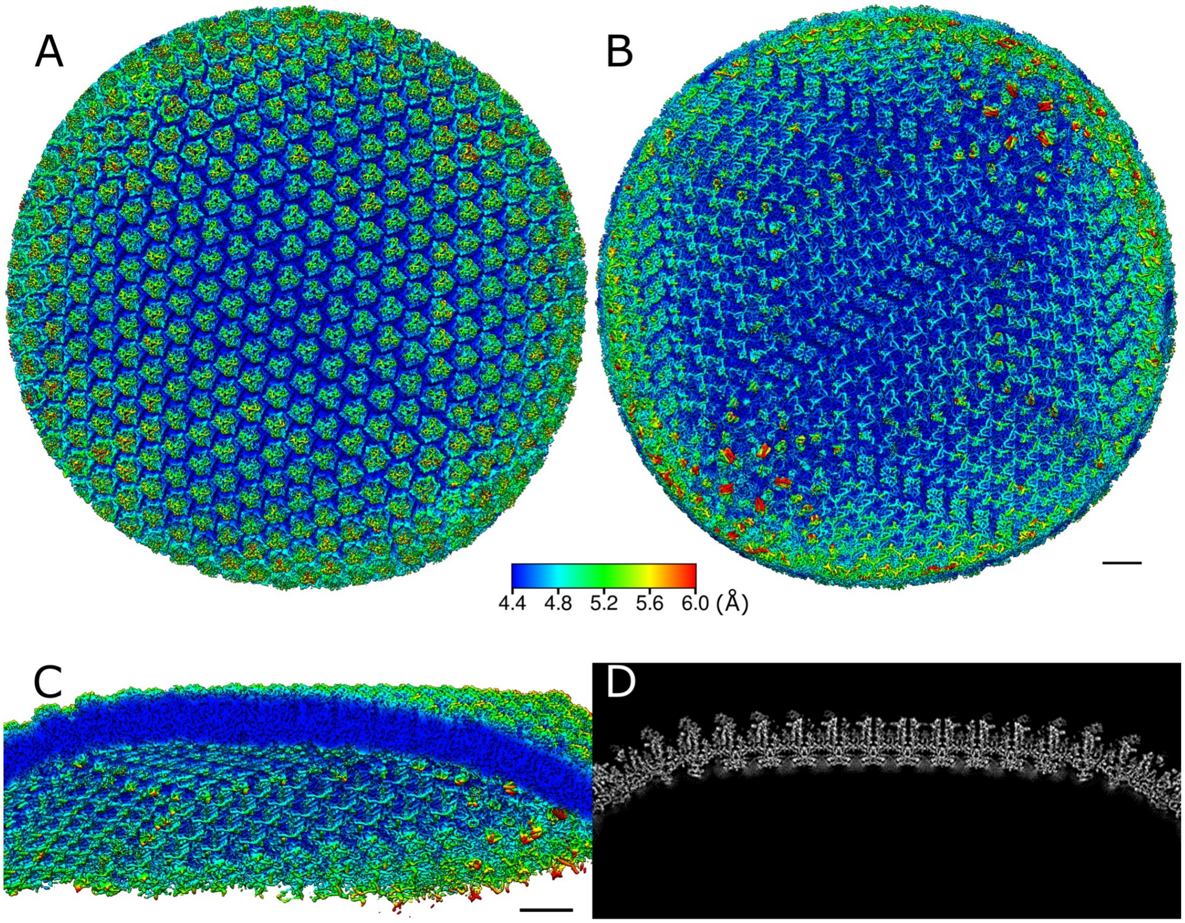


**Figure S5.** Block-based reconstruction of a two-fold axis point. A) External view. B) Internal view. C) Slice view perpendicular to (A) and (B). The map is colour coded according to the local resolution of the area. The colour code is shown in the figure. D) Slice view of the density map. Scale bars equal 10 nm.

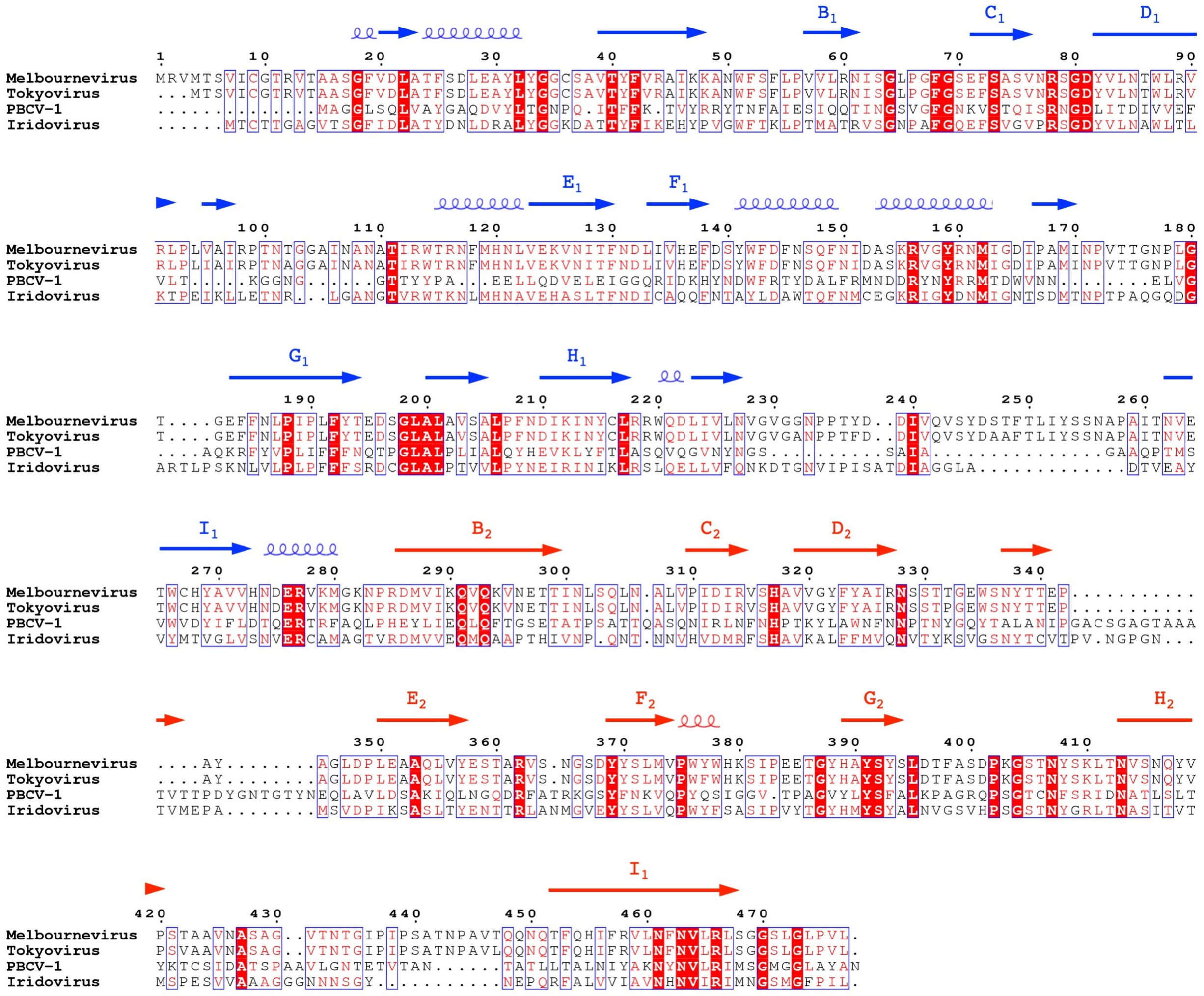


**Figure S6.** An alignment of Melbournevirus MCP against three other NCLDV MCPs, highlighting secondary structures: it is a helix with a coil and a β-sheet with an arrow. PROMALS3D (Pei et al., 2008) and ESpript3 (Robert and Gouet, 2014) were used. Labelled blue and red arrows indicate areas of β-sheet comprising the first and second jelly roll fold, respectively.

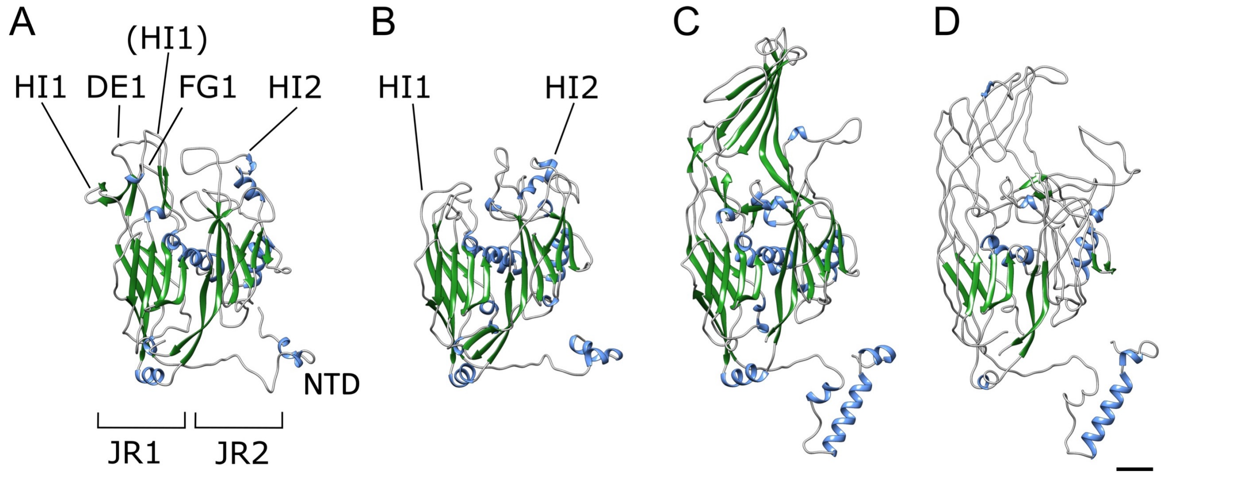


**Figure S7.** Comparison of the Melbournevirus (A), PBCV-1 (B) (Fang et al., 2019), faustovirus (C) (Klose et al., 2016) and ASFV (D) (Wang et al., 2019) MCP monomers, coloured by secondary structure. Random coil (grey), α-helix (blue) and β-sheet (green). The names of individual loops are indicated. JR1 and JR2 show jelly roll motif 1 and 2, respectively. NTD represents the N-terminal domain. Scale bar equals 1 nm.

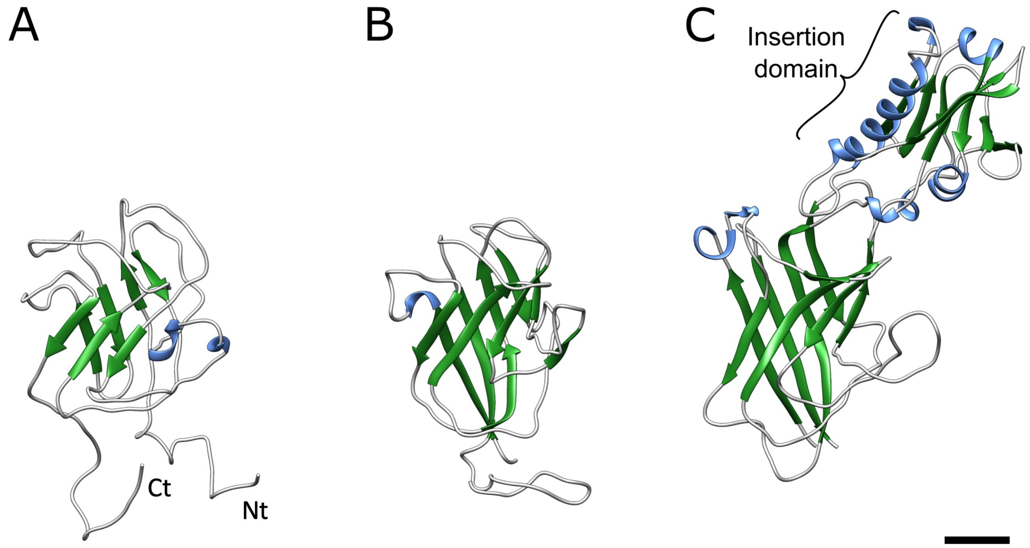


**Figure S8.** Comparison of (A) the ribbon model of the Melbournevirus penton base protein to (B) that of the PBCV-1 penton base protein (Fang et al., 2019) and (C) that of the CroV-dependent mavirus penton base protein (Born et al., 2018) with the single jelly roll motifs aligned. Ct and Nt show the C-terminal and N-terminal, respectively. Scale bar equals 1 nm.

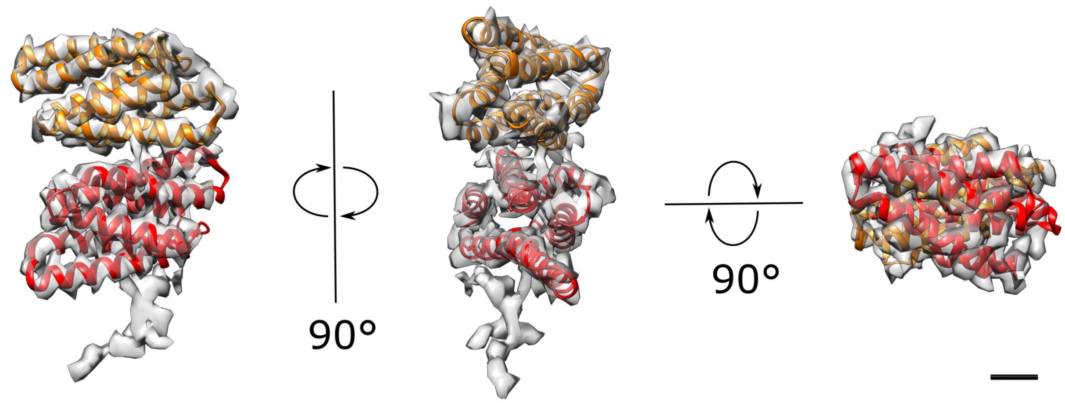


**Figure S9.** PC-α of Melbournevirus, with a *de novo* fitted polyalanine model. The mCP consists of two quadruple bundles of α-helices, stacked one on top of the other. Scale bar equals 1 nm.

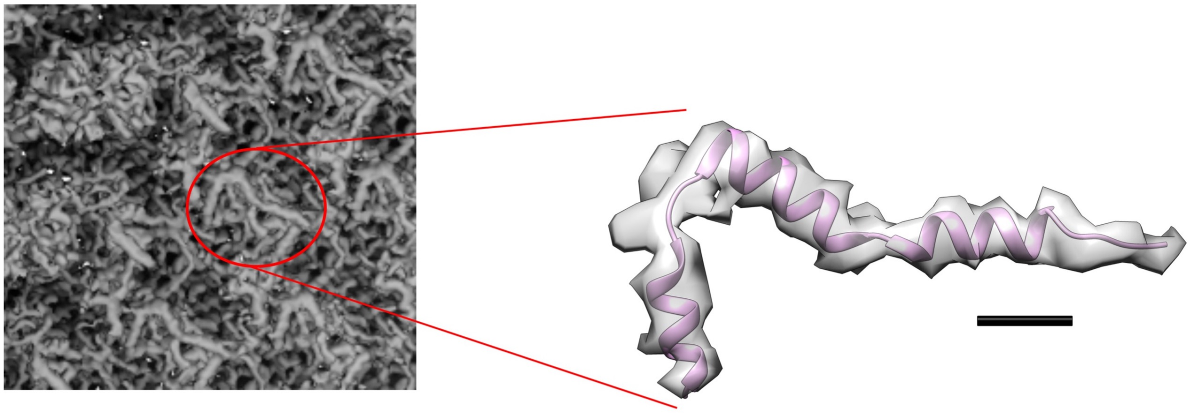

**Figure S10.** Lattice protein polyalanine model. Left, a close-up view of a section of the lattice from the two-fold block reconstruction, next to a zipper protein. Scale bar equals 1 nm.

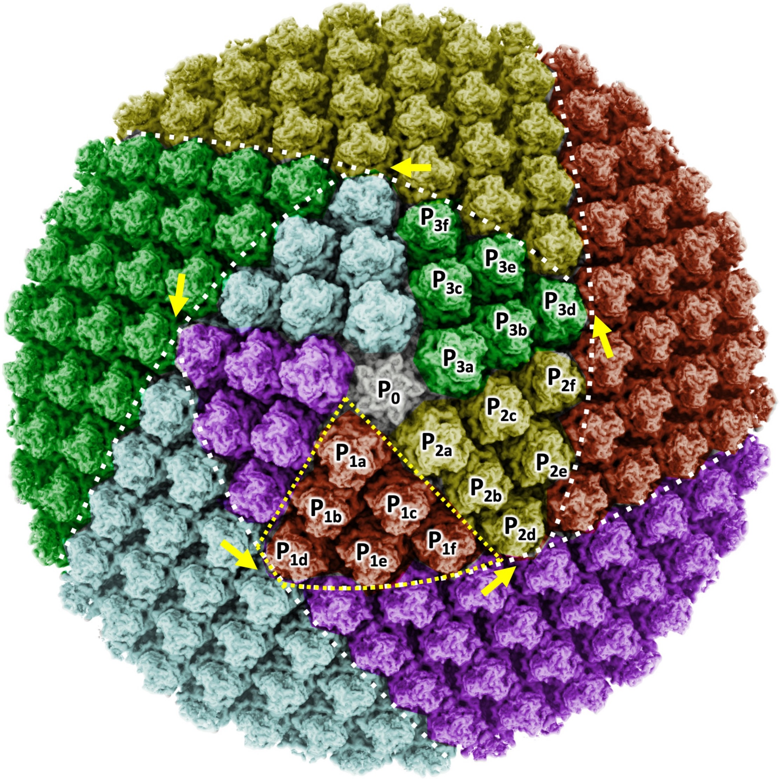


**Figure S11.** Alternate colouring of Fig. 5, if all the MCP trimers of the pentasymmetron asymmetric unit were oriented the same. The while dotted lines show the interfaces of trisymmetron and pentasymmetron. The yellow dotted trapezoid shows an asymmetric unit of the pentasymmetron. Three asymmetric units of the pentasymmetron are labelled $P_{1a\text{-}f}$, $P_{2a\text{-}f}$ and $P_{3a\text{-}f}$, showing that one of the MCPs in the asymmetric unit ($P_{1\text{-}3d}$) is not rotated 60°. The single MCP trimer would cause a mismatch in the MCP orientational alignment of the trisymmetron interface (yellow arrows). This shows how the smooth interface along the edge of the trisymmetron would be broken, potentially disrupting capsid formation.

### References

Born, D., Reuter, L., Mersdorf, U., Mueller, M., Fischer, M.G., Meinhart, A., and Reinstein, J. (2018). Capsid protein structure, self-assembly, and processing reveal morphogenesis of the marine virophage mavirus. Proc Natl Acad Sci U S A *115*, 7332-7337.

Fang, Q., Zhu, D., Agarkova, I., Adhikari, J., Klose, T., Liu, Y., Chen, Z., Sun, Y., Gross, M.L., Van Etten, J.L.*, et al.* (2019). Near-atomic structure of a giant virus. Nat Commun *10*, 388.

Fernandez-Leiro, R., and Scheres, S.H.W. (2017). A pipeline approach to single-particle processing in RELION. Acta Crystallogr D Struct Biol *73*, 496-502.

Klose, T., Reteno, D.G., Benamar, S., Hollerbach, A., Colson, P., La Scola, B., and Rossmann, M.G. (2016). Structure of faustovirus, a large dsDNA virus. Proc Natl Acad Sci U S A *113*, 6206-6211.

Pei, J., Kim, B.H., and Grishin, N.V. (2008). PROMALS3D: a tool for multiple protein sequence and structure alignments. Nucleic Acids Res *36*, 2295-2300.

Scheres, S.H.W. (2012). RELION: Implementation of a Bayesian approach to cryo-EM structure determination. J Struct Biol *180*, 519-530.

Robert, X. and Gouet, P. (2014) Deciphering key features in protein structures with the new ENDscript server. Nucleic Acids Res *42*, 320-324.

Wang, N., Zhao, D., Wang, J., Zhang, Y., Wang, M., Gao, Y., Li, F., Wang, J., Bu, Z., Rao, Z.*, et al.* (2019). Architecture of African swine fever virus and implications for viral assembly. Science *366*, 640-644.

Zivanov, J., Nakane, T., Forsberg, B.O., Kimanius, D., Hagen, W.J., Lindahl, E., and Scheres, S.H. (2018). New tools for automated high-resolution cryo-EM structure determination in RELION-3. Elife *7*, e42166.

Zivanov, J., Nakane, T., and Scheres, S.H.W. (2020). Estimation of high-order aberrations and anisotropic magnification from cryo-EM data sets in RELION-3.1. IUCrJ *7*, 253-267.
